# High temperature limits leaf size via direct control of cell cycle by coordinated functions of PIF4 and TCP4

**DOI:** 10.1101/2022.02.02.478810

**Authors:** Kumud Saini, Aditi Dwivedi, Aashish Ranjan

## Abstract

High ambient temperature suppresses Arabidopsis rosette leaf area, in contrast to elongation of stem and petiole. While the mechanism underlying temperature-induced elongation response is extensively studied, the genetic basis of temperature-regulation of leaf size is largely unknown. Here we show that warm temperature inhibits cell proliferation in the Arabidopsis leaves resulting in fewer cells compared to the control condition. Cellular phenotyping, genetic, and biochemical analyses established the key roles of PIF4 and TCP4 transcription factors in the suppression of Arabidopsis leaf area under high temperature by a reduction in cell number. We show that temperature-mediated suppression of cell proliferation requires PHYTOCHROME INTERACTING FACTOR 4 (PIF4). PIF4 interacts with TEOSINTE BRANCHED1/CYCLOIDEA/PCF4 (TCP4) and regulates the expression of cell cycle inhibitor, *KIP-RELATED PROTEIN 1* (*KRP1)*, to control the leaf size under high temperature. Warm temperature induces binding of both PIF4 and TCP4 to the *KRP1* promoter. PIF4 binding to KRP1 under high temperature is TCP4 dependent as TCP4 regulates PIF4 levels under high temperature. We propose a model where a warm temperature-mediated accumulation of PIF4 in the leaf cells promotes its binding to the *KRP1* promoter in a TCP4-dependent way to regulate cell production and leaf size. Our finding of high temperature-mediated transcriptional upregulation of *KRP1* integrates a developmental signal with an environmental signal that converges on a basal cell regulatory process.

**One sentence summary:** High ambient temperature suppresses Arabidopsis leaf area by reducing cell proliferation via PIF4- and TCP4-dependent transcription control of *KRP1*.

## Introduction

A suite of morphological adaptations that occurs in response to high ambient temperature is termed thermomorphogenesis, which includes hypocotyl and petiole elongation, accelerated flowering, etc. (Quint et al., 2016). While hypocotyl elongation is predicted to play a role in protecting meristematic tissue from the heat-absorbing soil, hyponastic petiole movement and open rosette structure help in increasing the leaf cooling capacity (Crawford et al., 2012; Kim et al., 2019). Leaves, on the other hand, show a reduction in blade area and thickness, reduced epidermal cell and stomatal density, early senescence, and reduced net carbon assimilation (Jin et al., 2011; Vile et al., 2012; Ibañez et al., 2017; Kim et al., 2020a). Leaves perceive and respond to various environmental signals, and are major above-ground mediators between a plant and its environment (Efroni et al., 2010; Lippmann et al., 2019). In the current warming climate, studying the impact of environmental changes on leaf growth and development and understanding the genetic mechanism for leaf phenotypic plasticity under varied environmental stimuli has become particularly important. While the mechanism underlying temperature-induced hypocotyl elongation is extensively studied, the genetic basis of temperature-regulation of leaf features is largely unknown (Gray et al., 1998; Koini et al., 2009; Quint et al., 2016). Leaves are lateral organs that arise as outgrowths from L1-L3 layers of shoot apical meristem. Recent reports show that the leaf is highly thermoresponsive and facilitates temperature-mediated growth in other aerial tissues (Kim et al., 2020b).

*PHYTOCHROME INTERACTING FACTOR 4* (*PIF4*) is a key transcription factor for thermomorphogenesis that integrates high-temperature cues to downstream phytohormone signaling and cellular response to manifest hypocotyl and petiole elongation (Koini et al., 2009; Quint et al., 2016). *PIF4* is expressed in all aerial tissue but its leaf epidermis-specific expression drives hypocotyl elongation and leaf hyponastic movements under high temperature (Kim et al., 2020b). In warm conditions, PIF4-mediated auxin induction in the leaves and ASYMMETRIC LEAVES 1 (AS1)- and PINOID (PID)- dependent directional auxin flow towards the abaxial side of the petiole determines the leaf thermonastic movements that help in leaf cooling (Park et al., 2019). PIF4-mediated transcriptional activation of growth-promoting genes, on the other hand, is antagonized by AT-hook transcription factors to suppress petiole elongation, suggesting a general involvement of PIF4 in growth regulatory responses (Favero et al., 2020). Recently, a role of PIF7 in temperature signaling was also reported in the early seedling growth and particularly for daytime regulation of high-temperature response, suggesting that different PIFs play specific roles in temperature-mediated growth adaptations (Chung et al., 2020; Fiorucci et al., 2020).

Several reports have suggested that PIF4 functions in concert with other growth regulators to control a variety of growth responses in a context-dependent fashion. For example, PIF4 regulates hypocotyl elongation in response to light and temperature via induction of auxin and brassinosteroids biosynthetic and response genes that involve integration with other transcription regulators, such as AUXIN RESPONSE FACTORs (ARFs), BRASSINAZOLE-RESISTANT 1 (BZR1), SEUSS, HOOKLESS 1 (HLS1), etc. (Huai et al., 2018; Ibañez et al., 2018; Reed et al., 2018; Jin et al., 2020). Apart from these transcription regulators, more recently, a small aromatic compound, Heatin, was shown to promote hypocotyl elongation via the PIF4-auxin module in response to temperature (Woude et al., 2021). Contrary to elongation response in hypocotyl and petioles, PIF4 is shown to suppress the formation of axillary meristems in dark along with BZR1 via inhibition of *ARGONAUTE10* and by antagonizing with ARF5 (Zhang et al., 2020). Temperature-mediated inhibition of meristemoid division via suppression of *SPEECHLESS* in the stomatal lineage cells is further reported to operate in a PIF4-dependent manner that leads to the production of fewer stomata in the cotyledons of Arabidopsis seedlings grown under high temperature (Lau et al., 2018). These reports suggest that PIF4 functions with other transcription factors in a context-dependent manner to play a pivotal role in linking environmental responses to the core developmental pathways. Thus, PIF4 crosstalk with other growth regulators needs to be explored for a better mechanistic understanding of these pathways and for their exploitation to optimize growth under a variable environment.

Five members of class II *TEOSINTE BRANCHED1/CYCLOIDEA/PCFs* (TCPs) regulated by miR319, control multiple developmental pathways including leaf growth, shape, and senescence (Palatnik et al., 2003; Schommer et al., 2012; Koyama et al., 2017). TCP4, one of the most characterized class II CIN-TCP transcription factors, suppresses cell proliferation by activating the transcription of a cell cycle inhibitor gene *KIP-RELATED PROTEIN 1* (*KRP1/ICK1*) and indirectly by repressing *GROWTH REGULATING FACTORs* (*GRFs*) via miR396 (Schommer et al., 2014). TCP4 was shown to promote cotyledon opening during the de-etiolation process by antagonizing the dark-induced PIF3, providing a molecular link between environmental signal and developmental response (Dong *et al*., 2019).

Plants display architectural plasticity in response to their surrounding environment via changes in organ shape or size by regulating basal cellular processes like cell division and expansion. Several reports have highlighted the importance of cell proliferation in mediating plant growth plasticity under various environmental stimuli (Tardieu and Granier, 2000; Rymen et al., 2007; Casadevall et al., 2013; Okello et al., 2016; Romanowski et al., 2021). This is achieved by tight regulation of cell cycle progression and arrest by various genes including several transcription factors, such as the NAC-type transcription factors ANAC044- and ANAC085-regulated cell cycle arrest under heat stress, *ICARUS1*-regulated cell proliferation under high temperature, miR396-mediated repression of GRFs to regulate cell proliferation under UV radiations, and more recently antagonistic regulation of cell cycle regulators by PIF7 and ANGUSTIFOLIA (AN3) to constraint leaf blade cell division by End-of-Day Far-Red (Casadevall et al., 2013; Zhu et al., 2015; Hussaini et al., 2021).

High temperature elicits different growth responses in different organs. Growth promotion by cell elongation is a well-documented response in plants under various environmental cues including high temperature. As opposed to hypocotyl and petiole elongation, high temperature-regulation of leaf size has not been studied in detail and the underlying genetic regulation is largely unknown. Here, we show that high temperature suppresses leaf area by inhibiting cell division in the leaves. We, further, show that temperature-induced PIF4 and TCP4 regulate the expression of *KRP1*, leading to inhibition of cell proliferation in leaves of the plants grown under high temperature.

## Results

### High-temperature-mediated suppression of leaf size involves inhibition of cell division

To investigate the effects of high ambient temperature on leaf morphological features, we sequentially examined the rosette area, individual rosette leaf area, epidermal cells, and palisade mesophyll cells of leaf four of the wildtype *Arabidopsis thaliana* ecotype, Col-0, under control temperature (21°C) and high ambient temperature (28°C). Col-0 rosette showed an open structure (i.e., increased convex hull) with a reduced total area under high temperature compared to the control (Figure 1A, Supplemental Figure 1A). Individual leaves also showed a reduction in their respective sizes at high temperature compared to the control (Figure 1B). Leaf four exhibited the most significant difference between the two temperature conditions among all the rosette leaves and, therefore, were used in this study for cellular phenotyping (Figure 1C, Supplemental Figure 1B). As expected, temperature-mediated changes in the lamina and petiole resulted in a decreased ratio of lamina and petiole length under high temperature (Supplemental Figure 1C). To find the cellular basis of leaf size reduction, we performed cellular profiling of leaf four by quantifying cell number and area in abaxial epidermal pavements cells and palisade mesophyll cells. We found a drastic 50% reduction in the number of epidermal cells per leaf and a 66% reduction in palisade cells under high temperature compared to the control (Figure 1D, Supplemental Figure 1D-F). In contrast, the average area of epidermal and palisade cells showed an increase under high temperature (Figure 1E, Supplemental Figure 1G). To confirm the results, we also quantified the epidermal cell number and area of the first pair of leaves under control and high temperature. The first pair of leaves also showed results similar to that of leaf four (Supplemental Figure 2). These results suggested that prolonged high temperature reduced leaf area by negatively impacting cell number that could likely result due to temperature-mediated inhibition of the cell division process.

**Figure 1.**
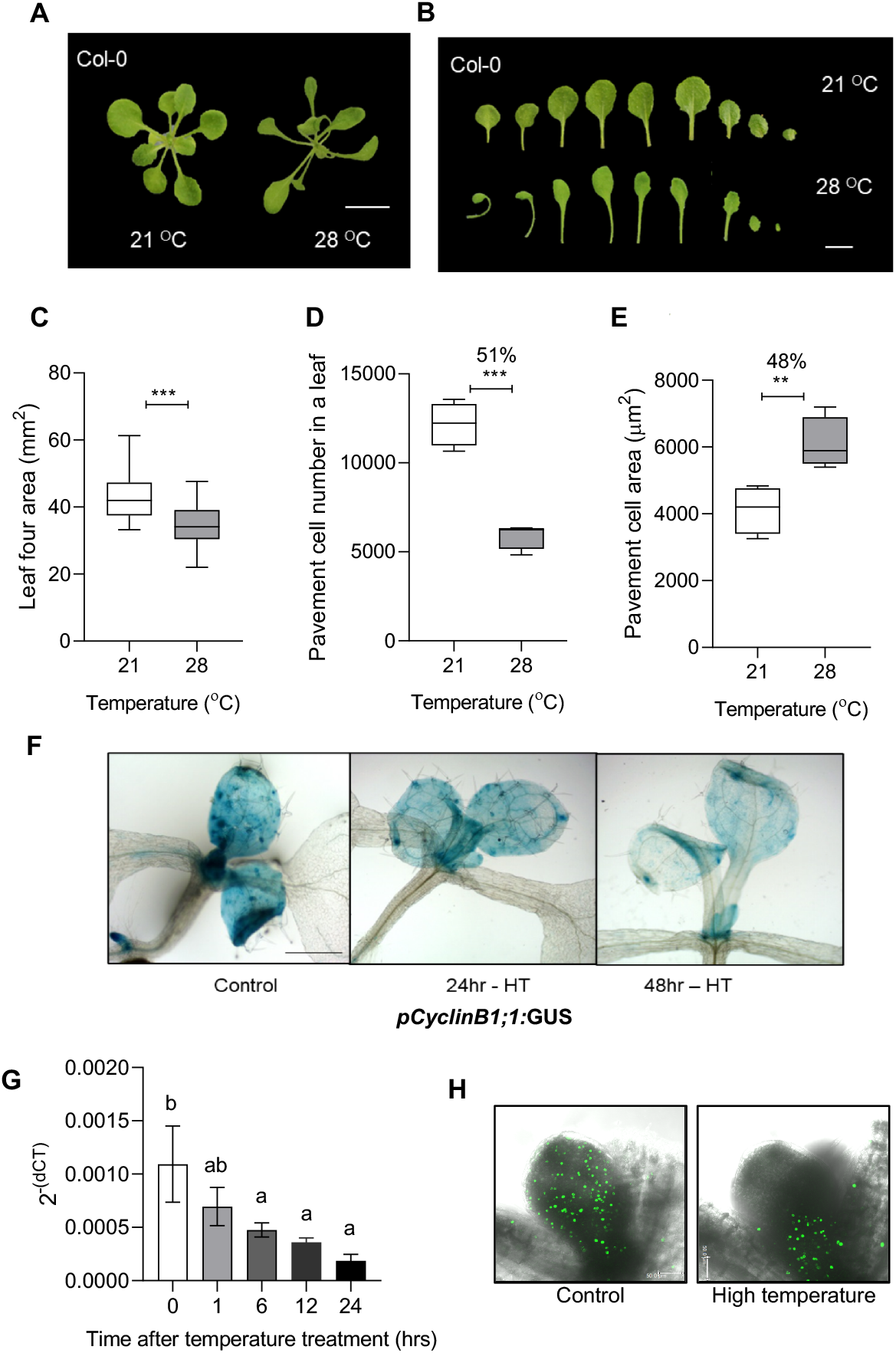
High temperature reduces leaf area by suppressing cell division in the Arabidopsis leaves. (A) Rosette picture and (B) leaf series of 21 ºC and 28 ºC grown Arabidopsis Col-0 wild type plants at 20 DAS. Scale, 1cm; DAS, Days after stratification. (C-E) Quantification of Leaf four area (n=19-23, C), Average pavement cell number on the abaxial side of leaf four (n=4, D), and Average pavement cell area (n=4, E) in 21 ºC and 28 ºC grown plant at 20 DAS. Box plot extends from 25th to 75th percentiles, where the line represents the median and whiskers minimum and maximum values. Percent change is indicated on top of the graphs. Stars indicate student’s t-test \*\*\**P* value <0.001. (F) GUS staining in *pCyclinB1;1:GUS* reporter line under control and high-temperature conditions. Scale, 1cm; HT - High Temperature. (G) Transcript levels of *CyclinB1;1* in the proliferating leaves in Col-0 at different time points after high-temperature treatment presented as 2^-(dCT)^ values. Values represent mean ± SE (n=3). Letters above the bars indicate statistically significant differences between the time points (one-way ANOVA followed by post hoc Tukey’s test, *P* < 0.05). (H) EdU fluorescence showing the distribution of mitotic cells in 5-day-old Col-0 leaf grown at control (21 ºC) and high temperature (28 ºC). Scale, 50 µm.

To test the effect of temperature on cell division, we first used a cell cycle marker line *pCyclinB1;1:GUS* (Donnelly et al., 1999). The marker line showed reduced GUS staining, and thus cell division, with increasing high-temperature exposure (Figure 1F, Supplemental Figure 3A). Consistent with *pCyclinB1;1:GUS* marker line, we detected reduced expression of *CyclinB1;1* in the proliferating leaves under high temperature (Figure 1G). We also detected reduced cell cycle progression using EdU (5-ethynyl-2’-deoxyuridine)-based S-phase assay in high temperature-grown seedlings compared to control (Figure 1H, Supplemental Figure 3B). Further, we used DAPI (4′,6-diamidino-2-phenylindole) to visualize mitotic figures in the basal region of young proliferating leaves as a readout for dividing cells and found a greater number of cells undergoing division under control condition than high temperature (Supplemental Figure 4). Taken together, these results showed that the high ambient temperature restricts the leaf size by inhibiting cell division.

### Temperature regulation of leaf size via cell division control is dependent on PIF4

The roles of PIF4 and PIF7 transcription factors in temperature-induced hypocotyl elongation response are well characterized (Koini et al., 2009; Chung et al., 2020; Fiorucci et al., 2020). However, the role of PIF transcription factors in temperature control of leaf growth is unexplored. Therefore, we first examined the expression levels of various PIFs in the leaves under high temperature. All the tested *PIF*s were upregulated shortly after high-temperature treatment, where *PIF4* showed the highest induction in its expression (Figure 2A). The induction in the expression, however, subsided at 8hrs of treatment for all the *PIFs* except *PIF4* (Supplemental Figure 5A). Consistent with this, the *pPIF4:GUS* line showed an increased spatio-temporal expression level after high-temperature treatment (Figure 2B, Supplemental Figure 5B). Interestingly, unlike the control leaves, high-temperature treated leaves showed increased GUS staining in the basal part of the leaf that coincides with the site of cell proliferation (Andriankaja et al., 2012). Using a translational reporter line, *pPIF4:PIF4-GFP*, we further showed comparatively more accumulation of PIF4 protein in the leaf epidermal cells under high temperature than the control (Figure 2C, Supplemental Figure 5C).

**Figure 2.**
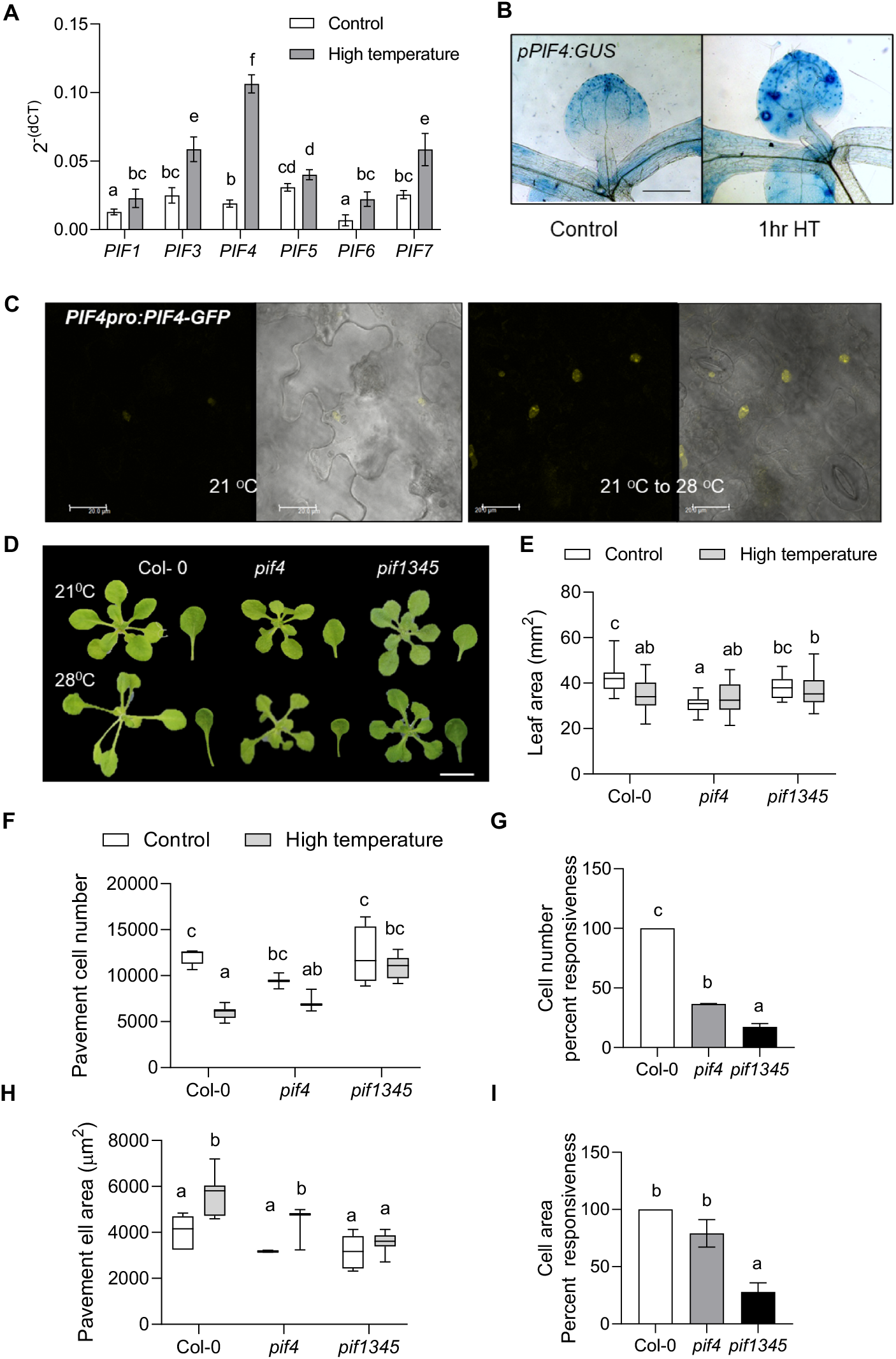
High temperature-mediated suppression of cell number involves PIF4. (A) Expression level of *PIF*s relative to an endogenous control, *ACTIN2*, in control and 1hr high-temperature treated young proliferating leaves of Col-0 plotted as 2-(dCT). Values represent mean ± SE (n=7-9). (B) GUS staining in control and high temperature treated leaves of *promoter:GUS* reporter lines of *PIF4*. HT - High Temperature. (C) Translation fusion reporter line *pPIF4:PIF4-GFP* showing protein localization in epidermal cells of control and high temperature treated seedlings Scale, 20µm. (D) Rosette and leaf pictures of different genotypes grown at 21 ºC and 28 ºC at 20 DAS. (E) Quantification of leaf four area (n=30-38). (F) Average pavement cell number in the abaxial side of leaf four (n=7-8). (G) Percent responsiveness of different genotypes to high temperature calculated with respect to the wild type Col-0 after setting the percent response of Col-0 to 100%. (H) Average pavement cell area in the abaxial side of leaf four (n=7-8). (I) Percent responsiveness of different genotypes to high temperature calculated with respect to the wild type Col-0 after setting the percent response of Col-0 to 100%. Box plot extends from 25th to 75th percentiles, where the line represents the median and whiskers minimum and maximum values. Different letters above the bars indicate statistically significant differences between genotypes or treatments (one-way ANOVA followed by post hoc Tukey’s test, *P* < 0.05).

Next, we investigated the leaf phenotypes of *pif4, pif7*, and, *pif1345* mutants under control and at high temperature. The rosette and leaf phenotype of *pif4* remain unchanged under high temperature compared to control condition, unlike Col-0 (Figure 2D, 2E, Supplemental Figure 5D & E). Quadruple mutant *pif1345* showed a decrease in rosette area, however, the area of leaf four remained unchanged under high temperature (Figure 2D, 2E, Supplemental Figure 5D). Notably, unlike Col-0, the cell number in *pif4* and *pif1345* mutants remained unchanged under high temperature (Figure 2F). To compare the different genotypes, we next calculated their responsiveness to high temperature-mediated cell number suppression with respect to Col-0. The percentage response in *pif4* was significantly reduced compared to a response in Col-0 which was set at 100%, suggesting involvement of PIF4 in cell number suppression (Figure 2G). It was further reduced in *pif1345*, suggesting that while PIF4 is sufficient for cell number reduction under high temperature, other PIFs could also play a role (Figure 2G). The epidermal cells in *pif4* showed expansion under high temperature, which could be explained by functions of other PIFs as evident from the *pif1345* mutant phenotype that failed to show cell expansion under high temperature (Figure 2H). Consistent with this, the responsiveness of pif*1345* towards cell expansion upon high-temperature induction was highly reduced compared to Col-0, while remained unchanged in *pif4* (Figure 2I). We found similar results for palisade cells that further proved PIF4 to be important for the regulation of cell proliferation in different layers of the leaf, while still being dispensable for cell expansion under warmth (Supplemental Figure 5F-I). Similar to leaf four, the average cell number per leaf in the first pair of leaves of the *pif4* mutant remained unchanged under high temperature (Supplemental Figure 2). Interestingly, *pif7*, unlike *pif4*, showed reduced rosette and leaf area and reduced cell number, however cell area remained unchanged (Supplemental Figure 6). Overall, these results suggested a direct and more important role of PIF4 in cell division control under high temperature possibly via changes in its spatio-temporal expression.

### TCP4 regulates leaf growth under high temperature

To get further insight on leaf size regulation under high temperature, we examined the publicly available transcriptome data to look for genes that expressed in the leaves and were also induced by high temperature (Stavang et al., 2009; Sidaway-lee et al., 2014). We then compared the list of temperature-induced genes with the list of literature curated leaf development-related genes (Supplemental File S1). Several genes encoding transcription factors, including several *TCPs* and *GRFs*, were found to be expressed in leaves as well as responsive to high temperature. We checked the transcript levels of temperature-responsive *TCPs* in leaves and found class II TCPs, *TCP3* and *TCP4*, to be highly upregulated by high temperature (Figure 3A). Consistent with this, a strong induction in GUS staining was visible in *pTCP4:GUS* reporter line within 1 hour of high-temperature treatment suggesting the potential involvement of *TCP4* in temperature signaling in the leaves (Figure 3B, Supplemental Figure 7).

**Figure 3.**
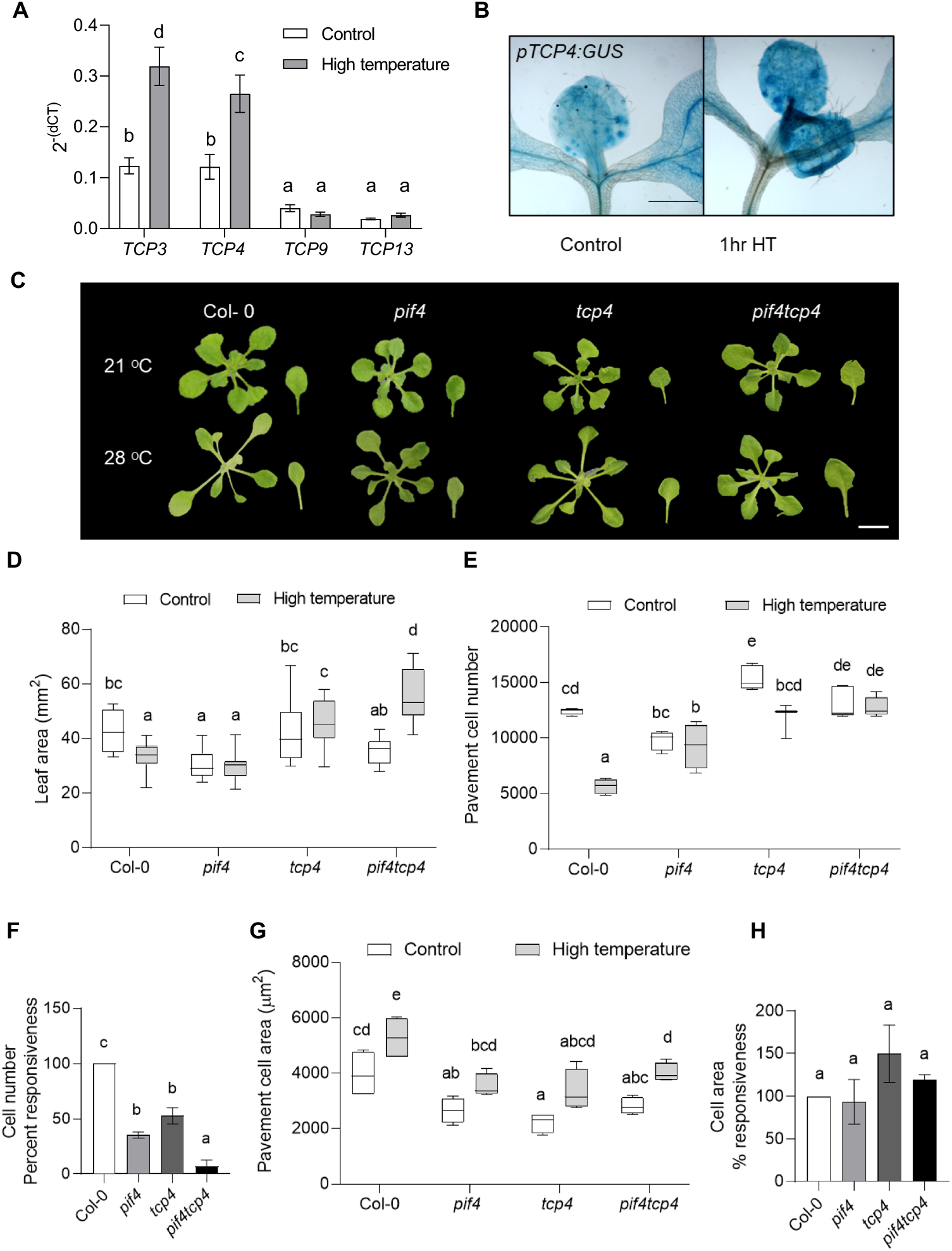
PIF4 and TCP4 co-regulate high temperature-induced cell proliferation inhibition. (A) Expression level of *TCP*s relative to an endogenous control, *ACTIN2*, in control and 1hr high-temperature treated young proliferating leaves of Col-0 plotted as 2-(dCT). Values represent mean ± SE (n=6-8). (B) GUS staining in control and high temperature treated leaves of *promoter:GUS* reporter lines of *TCP4*. HT - High Temperature. (C) Rosette and leaf pictures of different genotypes grown at 21 ºC and 28 ºC at 20 DAS. (D) Quantification of leaf four area (n=25) at 20 DAS in control and high temperature conditions. (E) Average pavement cell number in the abaxial side of leaf four (n=4). (F) Percent responsiveness of different genotypes to high temperature calculate with respect to the wild type Col-0 after setting the percent response of Col-0 to 100%. (G) Average pavement cell area in the abaxial side of leaf four (n=4). (H) Percent responsiveness of different genotypes to high temperature calculated with respect to the wild type Col-0 after setting the percent response of Col-0 to 100%. Box plot extends from 25th to 75th percentiles, where the line represents the median and whiskers minimum and maximum values. Different letters above the bars indicate statistically significant differences between genotypes or treatments (one-way ANOVA followed by post hoc Tukey’s test, *P* < 0.05).

To confirm the role of TCP4 in temperature-regulated leaf size control, we investigated the phenotype of the *tcp4* mutant. The leaf area remained unchanged in *tcp4*, however, the mutant showed an open rosette structure like Col-0 (Figure 3C, 3D, Supplemental Figure 8). Cell number reduced under high temperature, however, to a lesser extent than Col-0. Thus, the temperature-mediated cell number reduction response was attenuated in *tcp4* similar to as seen in *pif4* when compared to Col-0 (Figure 3E, 3F). Expansion in the epidermal cells in *tcp4* was comparable to Col-0 (Figure 3G, 3H). These results suggested that TCP4 is involved in temperature control of cell division in leaves but might not be for temperature-mediated petiole or cell elongation. To check the role of other members of miR319-regulated class II TCPs in temperature-mediated leaf size regulation, we used miR319 gain of function line, *jaw-1D* (*JAGGED AND WAVY-1D*), with multiple class II TCPs downregulated. *jaw-1D* mutant line displayed no change in cell number, cell area, or final leaf size between control and high temperature grown plants suggesting their role in temperature response (Supplemental Figure 6). A further attenuated cell number reduction response in *jaw-1D* than the single *tcp4* mutant suggested the involvement of multiple class II TCPs in temperature-controlled leaf growth regulation (Supplemental Figure 6F). *jaw-1D* rosette area, however, showed reduction under high temperature due to excessive leaf curling (Supplemental Figure 6A, B). These results established a role of Class II TCPs in temperature-regulated cell division control to determine leaf size.

### Both PIF4 and TCP4 are required for temperature-mediated regulation of cell number in the leaves

The expression levels of both *PIF4* and *TCP4* were induced under high temperature, and *pif4* and *tcp4* mutants showed unchanged leaf area with similar attenuation of cell division reduction under high temperature (Figure 3D-F). Based on these results, we hypothesized that the two transcription factors may function in a common signaling pathway, perhaps at the same time, to regulate leaf size under high temperature. To test this, we generated a *pif4tcp4* mutant line and it showed no change in rosette area but increased leaf four area under high temperature (Figure 3D, Supplemental Figure 8). Interestingly, *pif4tcp4* did not show any significant change in their cell number under high temperature and was irresponsive to the temperature-induced cell number suppression compared to Col-0 (Figure 3E, 3F). Partial attenuation of temperature-induced cell reduction response in *pif4* and *tcp4* single mutants and complete attenuation of the same in *pif4tcp4* double mutant demonstrated a direct involvement of both PIF4 and TCP4 in a common pathway that regulates cell number under high temperature. Unlike cell number changes, the cell expansion responses were similar to Col-0 (Figure 3G, 3H).

### PIF4- and TCP4- mediated cell cycle inhibition under high temperature occurs via *KRP1*

Since we hypothesized that PIF4 and TCP4 function in the same signaling pathway to control the temperature-mediated leaf size regulation, we intrigued about the signaling component downstream to PIF4 and TCP4. TCP4 is known to inhibit cell division via the upregulation of *KRP1* (Schommer et al., 2014). Hence, we first asked whether KRP1 could be involved in temperature-mediated cell cycle inhibition. The expression of *KRP1* was induced within 1hr of high-temperature treatment in Col-0. *pif4, tcp4*, and *pif4tcp4*, in contrast, did not show any change in *KRP1* expression under high temperature (Figure 4A). Interestingly, *KRP1* transcript levels were low in *pif4tcp4* compared to Col-0. These results suggested that temperature regulates *KRP1* expression and *KRP1* may function downstream to PIF4 and TCP4 to inhibit cell division.

**Figure 4.**
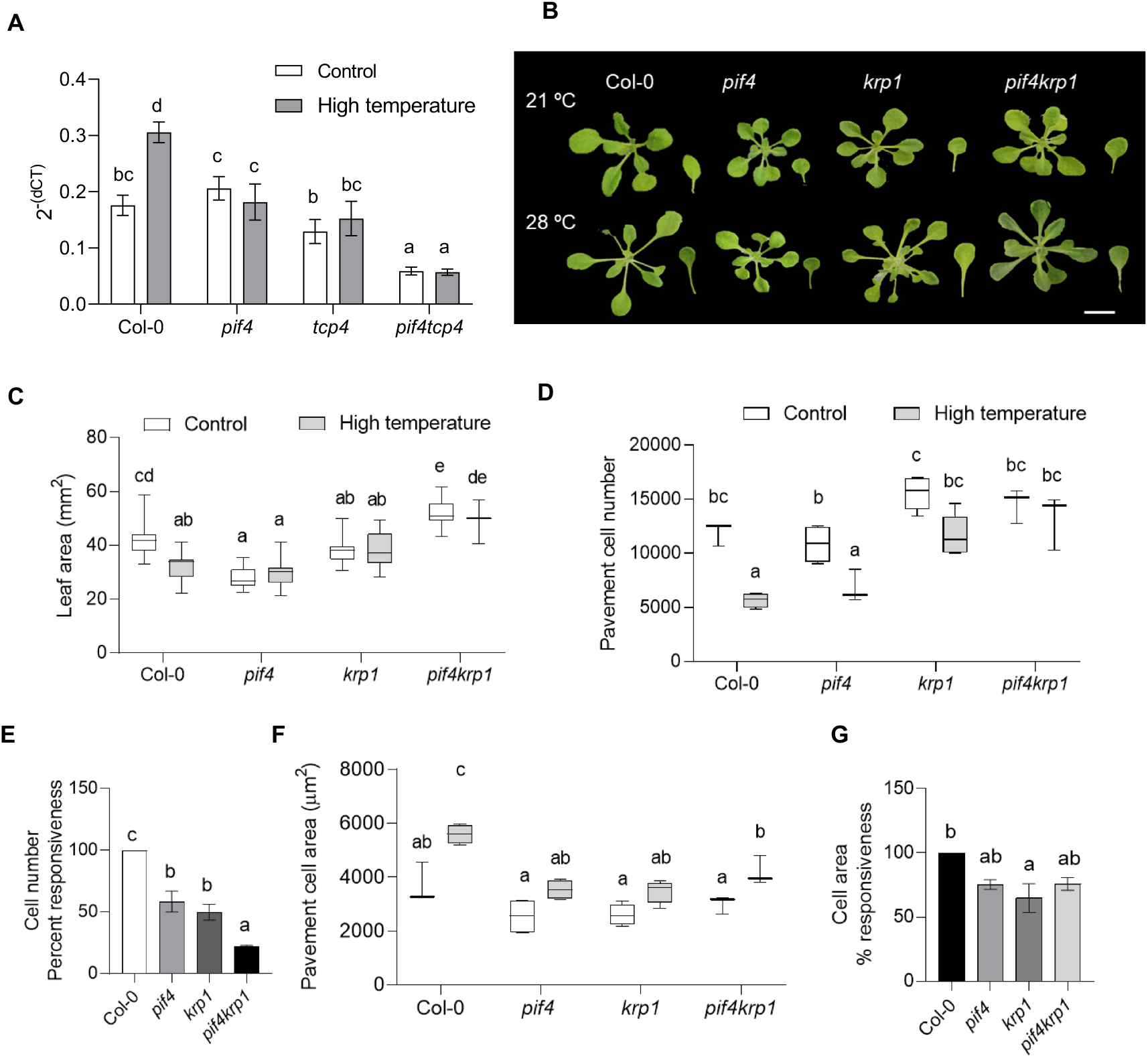
*KRP1* functions downstream of PIF4 to regulate cell number in leaf under high temperature. (A) Expression level of *KRP1* relative to an endogenous control, *ACTIN2*, in control and 1hr high-temperature treated young proliferating leaves in different genotypes plotted as 2^-(dCT)^. Values represent mean ± SE (n=6). (B) Representative rosette and leaf four images of 21 ^0^C and 28 ^0^C grown plants of Col-0, *pif4, krp1*, and *pif4krp1* at 20 DAS. (C) Quantification of leaf four area at 20DAS (n=8-12). (D) Average pavement cell number in the abaxial side of leaf four (n=3-5). (E) Percent responsiveness of different genotypes to high temperature calculate with respect to the wild type Col-0 after setting the percent response of Col-0 to 100%. (F) Average pavement cell area in the abaxial side of leaf four (n=3-5). (G) Percent responsiveness of different genotypes to high temperature calculated with respect to the wild type Col-0 after setting the percent response of Col-0 to 100%. Box plot extends from 25th to 75th percentiles, where the line represents the median and whiskers minimum and maximum values. Different letters above the bars indicate statistically significant differences between genotypes or treatments (one-way ANOVA followed by post hoc Tukey’s test, *P* < 0.05).

To investigate the involvement of KRP1 in leaf size regulation under warm temperature, we looked for the leaf phenotype of the *krp1* mutant. Rosette and leaf area and cell number and area showed no significant change under temperature treatment (Figure 4C-F, Supplemental Figure 9). Similar to *pif4, krp1* showed attenuated response for cell number reduction compared to Col-0 under high temperature (Figure 4E). To test for functional redundancy among the *KRP*s, we checked the phenotype of the triple mutant *krp4/6/7*. The triple mutant showed a smaller rosette and leaf with a significant reduction in their cell number under high temperature, similar to Col-0, excluding the possibility of their involvement in temperature-regulated leaf size control (Supplemental Figure 6C-G). To test whether *PIF4* and *KRP1* work in the same pathway to control leaf size under high temperature we examined the phenotype of the *pif4krp1* double mutant. Rosette and leaf area were unchanged between the control and high-temperature grown plants (Figure 4B 4C, Supplemental Figure 9). As hypothesized, cell numbers were unaffected in *pif4krp1* by high-temperature treatment along with further reduced responsiveness of the double mutant than either of the single mutant when compared to Col-0 (Figure 4D, E). Cell area increased in *pif4krp1* similar to Col-0 (Figure 4F, 4G). Overall, these results supported our hypothesis that PIF4 and KRP1 may act in the same pathway to control cell number in high-temperature grown leaves.

### PIF4 and TCP4 coordinate to regulate *KRP1* expression

TCP4 is known to directly activate the expression of *KRP1*, we checked whether PIF4 regulates *KRP1* expression (Schommer et al., 2014). Using luciferase transactivation assay, we showed that PIF4 induced *KRP1* expression (Figure 5A, 5C). We also validated the binding of TCP4 to the *KRP1* promoter to induce the expression (Figure 5B, 5C). In addition, we quantified the relative activity of the *KRP1* promoter using dual luciferase assay. We found more *KRP1* promoter activity in presence of both PIF4 and TCP4 compared to the individual transcription factors (Figure 5C). This ruled out competitive binding between the two transcription factors on the *KRP1* promoter and suggested that both could potentially co-regulate *KRP1* expression. This further raised the possibility of physical interaction between PIF4 and TCP4 to co-regulate the downstream target(s). We, therefore, performed a protein-protein interaction assay using a split luciferase assay and BiFC in *Nicotiana benthamiana* leaves and showed a positive interaction between PIF4 and TCP4 (Figure 5D, 5E). We further proved this interaction in a heterologous system using a yeast two-hybrid assay (Figure 5F). Since the temperature-mediated reduction in cell number was completely abolished in the *jaw-1D* mutant, it is plausible that PIF4 might interact with other members of Class II TCPs. Protein-protein interaction study using BiFC showed that indeed PIF4 interacted with TCP2, TCP3, TCP10, and TCP24 apart from TCP4 in that class (Supplemental Figure 10).

**Figure 5.**
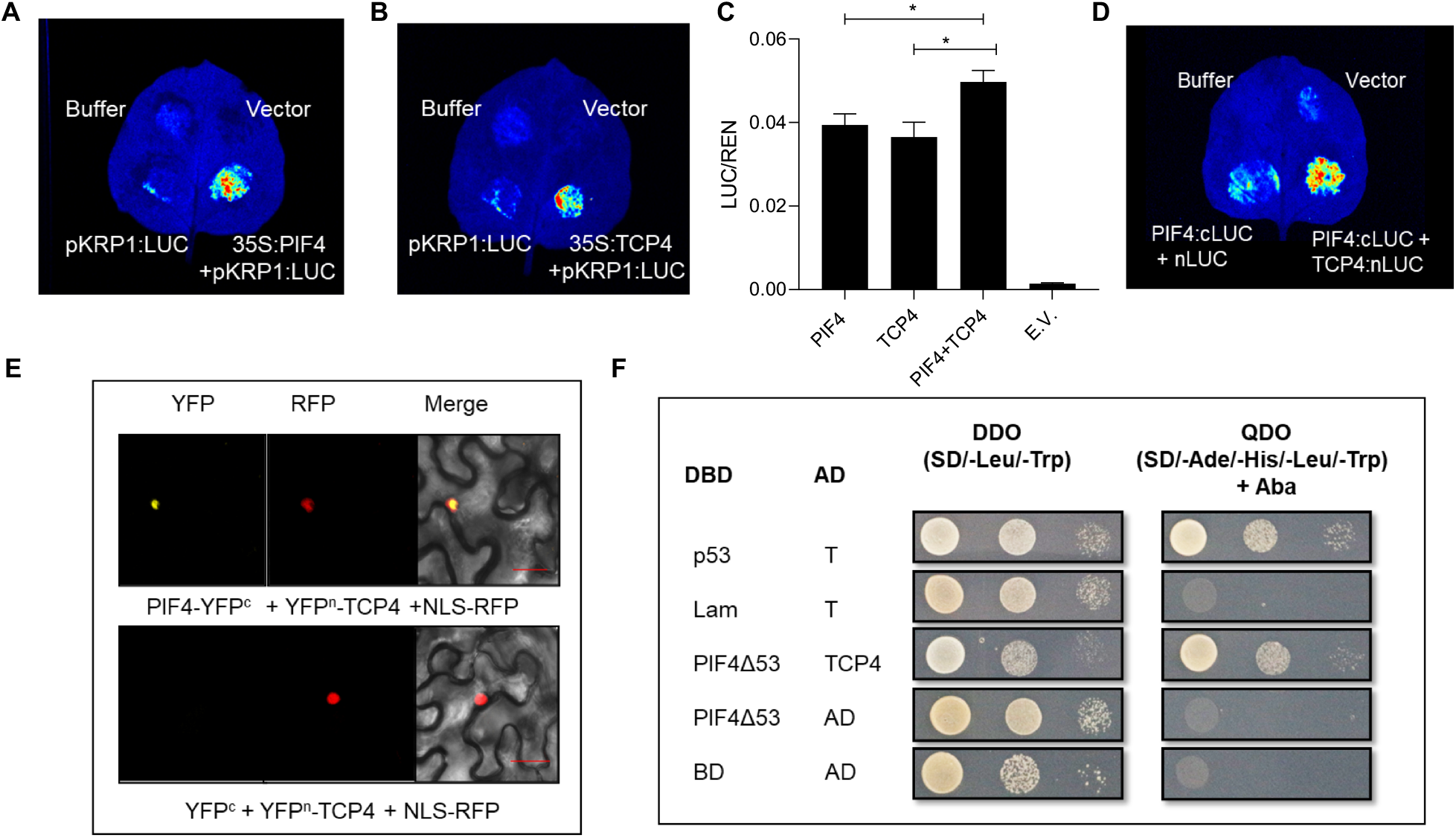
PIF4 interacts with TCP4 and regulates *KRP1* expression. (A, B) Luciferase transactivation assay showing binding of PIF4 (A) and TCP4 (B) to the *KRP1* promoter in *Nicotiana benthamiana* leaves. (C) Dual-luciferase assay in *Nicotiana benthamiana* leaves co-infiltrated with different transcription factors. LUC/REN ratio represents the relative activity of the *KRP1* promoter, where REN was used as an internal control. Values represent mean ± SE (n=5) E.V.= empty vector. Stars indicate student’s t-test *P-*value <0.05. (D) Split Luciferase assay showing protein-protein interaction between PIF4 (cloned with cLUC) and TCP4 (cloned with nLUC). (E) BiFC assays in *N. benthamiana* showing an interaction between PIF4 and TCP4. NLS-RFP is used as a nuclear marker. Scale, 30 µm. (F) Yeast two-hybrid assays showing an interaction between PIF4Δ53 and TCP4. The DNA-binding domain (DBD) was fused to truncated PIF4 named as PIF4Δ53, and the activation domain (AD) was fused to TCP4.

To get further insight into the co-regulatory mechanism of *KRP1* by PIF4 and TCP4, we scanned the promoter of *KRP1* for putative PIF4 binding sites. Within 1 kb upstream of the start codon, we found two putative PIF4-binding G-boxes close to the known TCP4 binding site on the *KRP1* promoter (Figure 6A). Using Chromatin Immunoprecipitation (ChIP)-PCR, we first confirmed the binding of TCP4 on the *KRP1* promoter and interestingly found it to increase by nearly forty-fold by exposure to high temperature (Figure 6B). Next, we did ChIP-qPCR using anti-PIF4 antibodies under control and high temperature-treated samples in different genotypes. We observed a four- and two-fold enrichment of putative G-box1 and G-box2 at *KRP1* promoter using anti-PIF4 antibody after high-temperature treatment in Col-0 (Figure 6C). Interestingly, we didn’t find increased enrichment of G-boxes on the promoter of *KRP1* in the *tcp4* mutant under high temperature, suggesting a requirement of TCP4 for PIF4 to regulate KRP1. We confirmed the TCP4-dependent PIF4 regulation of gene expression using *IAA29*, which is an auxin-responsive gene known to be thermo-responsive in a PIF4-dependent manner, as a positive control (Figure 6D). Since TCP4 is a transcription factor, the regulation of *PIF4* expression by TCP4 would be the most plausible mechanism for this. We detected increased enrichment of the two putative TCP binding elements on the PIF4 promoter under high temperature (Figure 7A). In support, *PIF4* expression failed to elicit in response to high temperature in *tcp4*, contrary to Col-0, suggesting transcriptional level regulation that further affected PIF4 protein abundance and thus PIF4 availability for binding to its targets (Figure 7B). Indeed, western blot results confirmed that TCP4-dependent regulation of PIF4 level under high temperature, as the mutant failed to show increased PIF4 levels under high temperature as seen in Col-0 (Figure 7C). Taken together, we showed a TCP4-dependent PIF4-mediated regulation of *KRP1* under high temperature that regulate leaf size in warmth (Fig. 7D).

**Figure 6.**
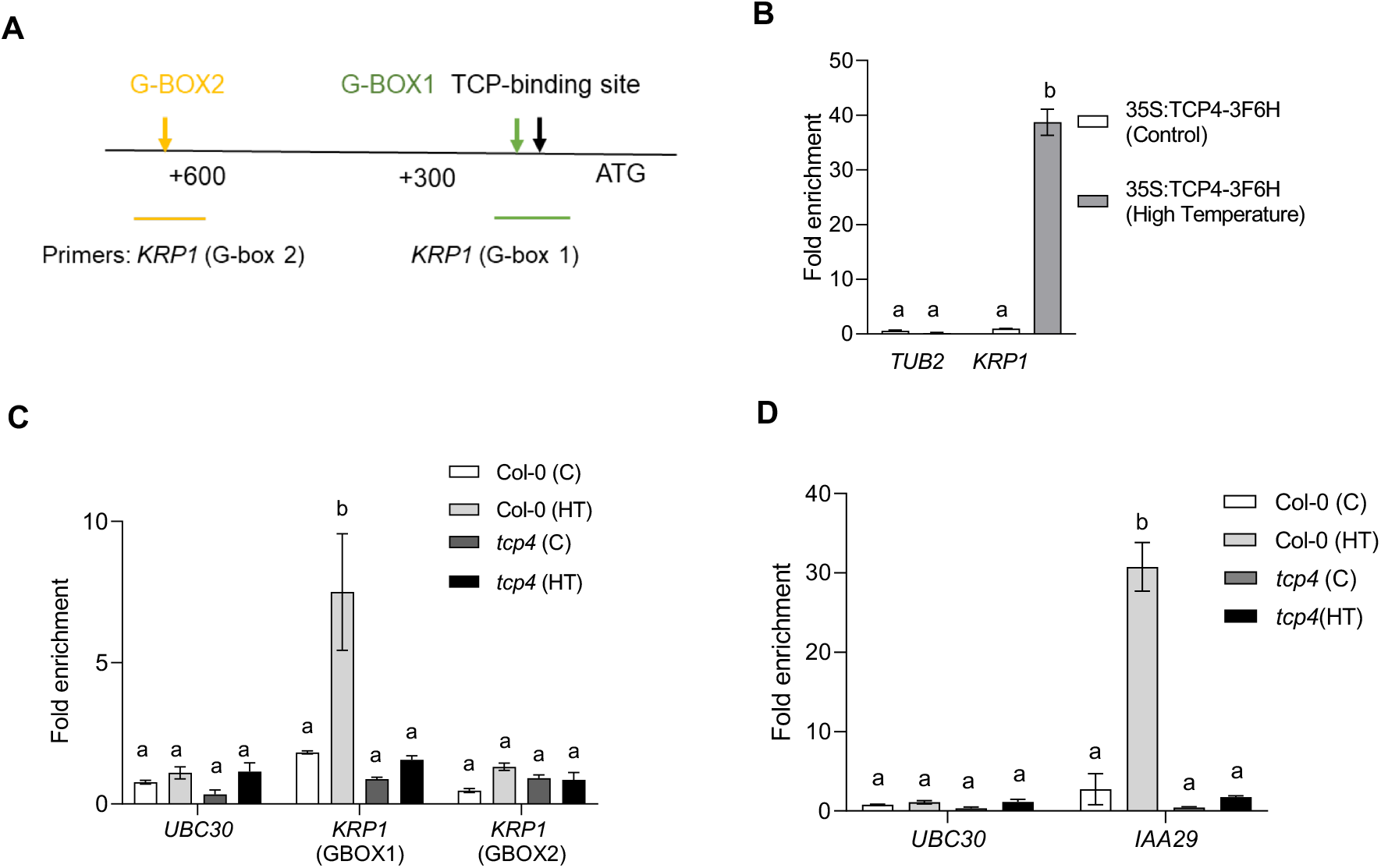
PIF4 regulates *KRP1* in a TCP4-dependent manner. (A) Schematic diagram of *KRP1* promoter with ATG start site showing the presence of TCP4 binding site and putative G-boxes. (B) Chromatin Immunoprecipitation (ChIP) assay showing enrichment of TCP-binding element in the promoter of *KRP1* region in *35S:TCP4-3F6H* line using anti-FLAG antibody under control and high temperature. (C)ChIP assay showing enrichment of G-box region using the anti-PIF4 antibody in the promoter of *KRP1* in Col-0 and *tcp4* under control and high temperature along with negative control *UBC30*. (D) Enrichment of G-box region using the anti-PIF4 antibody in the promoter of *IAA29* under control and high temperature in Col-0 and *tcp4* along with negative control *UBC30*. Values represent mean ± SE (n=3-4). Different letters above the bars indicate statistically significant differences between genotypes or treatments (one-way ANOVA followed by post hoc Tukey’s test, *P* < 0.05). C-Control, HT-High Temperature

**Figure 7.**
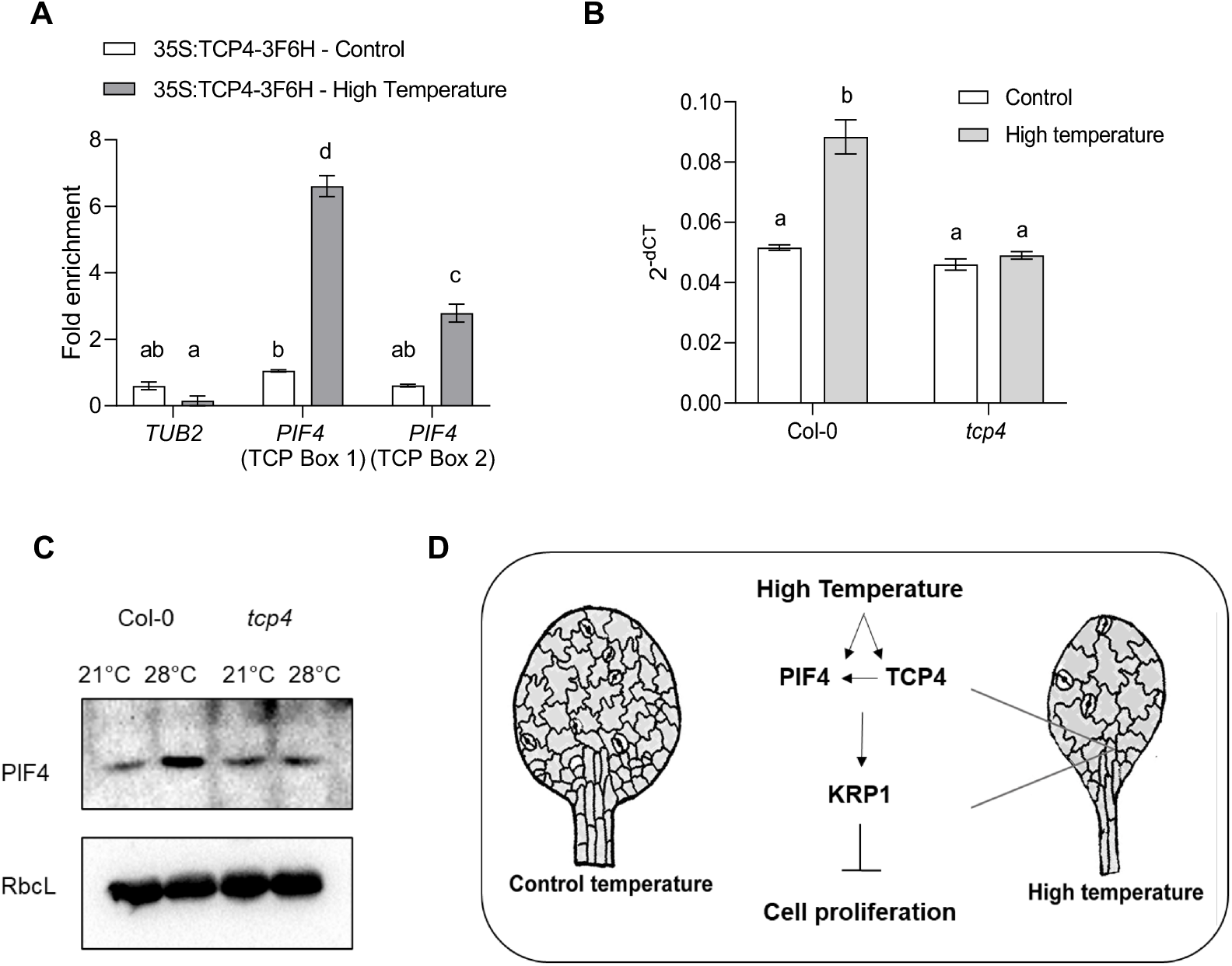
TCP4 regulates temperature-induced PIF4 levels. (A) ChIP assay showing enrichment of TCP binding sites on the promoter of *PIF4* under control and high temperature in *35S:TCP4-3F6H* line using anti-FLAG antibody. (B) Quantification of *PIF4* transcript level in proliferating leaves of Col-0 and *tcp4* under control and 3-hour temperature-treatment represented as 2^-(dCT)^. Values represent mean ± SE (n=4). Letters above the bars indicate statistically significant differences between genotypes or treatments (one way ANOVA followed by post hoc Tukey’s test, *P* <0.05). (C) Western blot comparison of PIF4 protein levels in Col-0 and *tcp4* background at 21 °C and after high temperature (28 °C) treatment between Zt1 to Zt5. RbcL was used as the loading control. (D) A schematic diagram representing high-temperature-mediated suppression of leaf size. Warm temperature-mediated accumulation of PIF4 in the leaf cells promotes its binding to the *KRP1* promoter in a TCP4-dependent way to reduce cell production and thus leaf size.

## Discussion

Plants show remarkable plasticity in their growth and development in response to their surroundings by tightly regulating the final organ size and shape. Shade avoidance syndrome is one of the well-studied classical examples of such growth plasticity where plants exhibit various architectural adaptions including the elongation response to compete with neighboring vegetation for light (Pierik and De Wit, 2014). Recently, the role of cell division in regulating leaf size under shade was highlighted, where shade-grown leaves showed fewer cells and thus small size as opposed to longer hypocotyl with elongated epidermal cells (Romanowski et al., 2021). Cell division and cell expansion are the key processes to mediate plant growth and developmental plasticity at the cellular level (Hepworth and Lenhard, 2014). Temperature-induced growth plasticity varies in different organs, and these organ-specific responses become evident at the early seedling development itself. While studies report a slight increase in the number of epidermal cells in the hypocotyl, the organ elongation response under various environmental cues including high temperature is largely attributed to the cell expansion process (Gray et al., 1998; Ibañez et al., 2018; Bellstaedt et al., 2019). However, at least one study reported a reduction in stomatal density in the high-temperature-grown cotyledons (Lau et al., 2018). Here we show a central role of cell division for leaf growth plasticity in response to temperature. We report that PIF4 and TCP4 play critical and co-dependent roles to regulate cell number in high-temperature grown leaves by suppressing the expression of *KRP1* (Figure 7D). Together, we showed that coordinated interplay between PIF4, an environmental regulator, and TCP4, a developmental regulator, determines the fate of final leaf size under high temperature.

### Temperature regulates leaf size via cell division control

Temperature-induced cell elongation response is shown to drive leaf thermonastic movement that assists in plant cooling via transpiration (van Zanten et al., 2010; Crawford et al., 2012). Smaller and thinner leaves, on the other hand, help in heat dissipation via evaporation and convection (Franklin and Wigge, 2014; Quint et al., 2016). While a 1-3 °C rise in temperature was found to be advantageous for Arabidopsis, a 5 °C or more rise is known to adversely affect leaf functional traits, including photosynthesis. Thus, the reallocation of reserved carbon in driving elongation growth could be a mean to increase cooling via transpiration or to reach a light source to increase photosynthetic capacity (Jin et al., 2011; Romero-Montepaone et al., 2021). In agreement with this, Vasseur et al. (2011) has shown that plants only exhibit temperature-induced elongation response under low to medium light. The effect of temperature on plant phenotype subsides when light is not limiting. This suggests that elongation response seen under high temperature could be primarily to increase light capture and photosynthetic capacity. However, as high temperature affects a wide array of changes in both vegetative and yield-related traits, this remains to be seen how those will vary under varying light and temperature conditions (Ibañez et al., 2017).

Previously, temperature-mediated hypocotyl elongation was shown to involve the expansion of epidermal cells (Gray et al., 1998). We studied the cellular and molecular basis of smaller leaf size under high temperature compared to the control temperature. Cellular profiling of leaves in combination with the expression pattern of cell cycle markers in response to high temperature confirmed that temperature-mediated restriction of leaf size involves suppression of cell division (Figure 1). Interestingly, a previous report showed enrichment of several cyclins, KRPs, and CDKs involved in cell cycle regulation among the genes differentially expressed in Arabidopsis plants grown under high temperature (Sidaway-Lee et al., 2014). Our results suggested possibilities of overall reduced cell division and/or changes in the duration of the cell division phase in the leaves under high temperature. Prolonged exposure to shade, showing plant responses similar to high temperature, has been reported to cause an early exit from cell proliferation, suggesting that high temperature and shade likely control the leaf size via control of cell division (Carabelli et al., 2018). Consistent with this, a recent report has suggested a Phytochrome B (PhyB)-dependent suppression of CYCB1;1 expression and cell proliferation in the leaves under shade (Romanowski et al., 2021). PhyB is reported to be a thermosensor in Arabidopsis, suggesting PhyB likely integrates high-temperature signal to repression of cell division response via downstream transcription factors, such as PIFs (Jung et al., 2016; Legris et al., 2016).

### PIF4 as a mediator of plant’s environment to growth plasticity

PIF4 expression domain is widespread throughout the leaf development as it accumulates in the epidermis of young leaves, at the SAM/Leaf boundary, in the middle zone of leaf coinciding with leaf vasculature framework, and later at the axils of the young leaves (Zhang et al., 2020). The spatio-temporal expression pattern of *PIF4* suggests a potential role of the gene at various stages of leaf and meristem patterning. Our results showed that PIF4 accumulates in the leaf epidermal cells under warm temperature and suppresses cell number, as was evident from the *pif4* mutant showing attenuation of cell number reduction under high temperature compared to Col-0 (Figure 2). The role of PIF4 in elongation response is extensively studied in the past decade (Koini et al., 2009; Choi and Oh, 2016; Ibañez et al., 2018). More recently, evidence for the involvement of PIF4 in the regulation of cell division is reported in the axillary meristem and stomatal lineage cells (Lau et al., 2018; Zhang et al., 2020). This suggests that PIF4 could potentially control the fate of the cells leaving the meristems for division or differentiation in the leaves. Whether it is the thermonastic movements to induce leaf cooling or reduction in leaf size to help in heat dissipation, both the responses involve differential regulation of PIF4 for cell elongation vs cell division. Together, these emphasize the pivotal importance of PIF4 in complex growth regulatory decisions that plants need to make under high temperature. Hussaini *et al*. (2021) showed PIF7 as a major player in limiting cell division in shade-grown plants via competing with AN3 that is known to promote cell proliferation along with GRFs. Interestingly, based on the cellular phenotype we did not find major involvement of PIF7 in cell number suppression rather in cell elongation response under high temperature (Supplemental Figure 6).

### TCP4 as a regulator of leaf growth plasticity in warm conditions

Class I and class II of TCP transcription factors antagonistically control growth. Class I members, in general, promote cell proliferation, while class II members promote cell differentiation and progression of cell cycle arrest front (Gutie et al., 2005; Efroni et al., 2008; Kieffer et al., 2011). Recent reports have shown the involvement of three class I TCPs in temperature signaling, where, TCP 5, 13, and 17 were shown to regulate as well as interact with PIF4 in temperature-induced hypocotyl elongation response (Han et al., 2019; Zhou et al., 2019). We observed a similar regulatory response between TCP4 and PIF4 in leaf size regulation in warm temperature. TCP4 was shown to promote CONSTANS levels to induce photoperiodic flowering (Kubota et al., 2017). However, the role of class II TCPs in the regulation of temperature response is unexplored. Our results established a critical role of TCP4 in temperature-induced leaf size reduction based on its reduced response to cell suppression under warm conditions. Although a non-significant change in cell number in *jaw-1D* suggests that downregulation of multiple class II TCPs could also mitigate the effect of temperature on cell division (Supplemental Figure 6). TCP4 is already known to inhibit cell division and promotes cell differentiation in leaves, our study adds a new dimension to TCP4 function in restriction of leaf size via cell division inhibition under high temperature.

### TCP4 regulates PIF4 while working alongside

Our cellular, genetic and biochemical analyses proved that TCP4 works alongside PIF4. We showed similar attenuation in cell number reduction under high temperature in *tcp4* and *pif4* mutants and a complete irresponsiveness to changes in cell number in *pif4tcp4* suggesting their critical and co-regulatory role in the cell division process (Figure 3). Cell expansion changes in *pif4* or *tcp4* leaves were comparable to Col-0 making them dispensable for the process. However, the response was reduced in *pif1345* and *jaw1-D* mutants suggesting redundancy in the gene families for cell expansion response. Similar to PIF4, TCP4 is also known to promote cell elongation via activation of genes involved in auxin and brassinosteroids biosynthesis and signaling (Challa *et al*., 2016). This suggests that TCP4, along with PIF4, could be instrumental in integrating environmental signals and developmental responses. A TCP4-dependent regulation of PIF4 levels and, in turn, PIF4 availability on its target binding sites, along with evidence of a physical protein-protein interaction between the two further highlights the complexity and degree of regulation of their targets. Collectively, our genetic analyses, as well as DNA-protein interactions, showed that PIF4 and TCP4 work in a common pathway to control cell proliferation, and possibly other aspects of leaf growth and development that have the potential to be explored further.

### PIF4 and TCP4 integrate high-temperature signal to cell cycle inhibitor KRP1

Previous studies have shown leaf area reduction via a reduction in cell number under mild osmotic stress, UV irradiation, and simulated shade suggesting that inhibition of cell cycle machinery could be a general response by which plants modulate their growth under diverse environmental conditions (González et al., 1998; Wargent et al., 2009; Skirycz et al., 2011). Leaf growth and development is a complex growth process, for simplicity and dissection of novel pathways under high temperature, we largely focused on temperature-mediated cell number suppression and leaf size regulation in plants grown under continuous high-temperature conditions post-germination. We provided a molecular link connecting the core leaf development pathway and high-temperature signaling via *KRP1. KRP1* showed upregulation under high temperature in a PIF4- and TCP4-dependent manner to regulate cell number and leaf size (Figure 4). Increased expression of *KRP1* negatively impacts cell division and plant growth by inhibiting the activity of cyclin-dependent kinases (CDKs) (Wang et al., 2000). Attenuated cell number suppression phenotype in the mutants and TCP4-dependent abundance of PIF4 on the *KRP1* promoter suggest that the three represent the core temperature signaling components that regulate leaf size (Figure 7D). In addition, the large-sized leaf phenotype in *pif4tcp4* suggests that the two transcription factors may be involved in pleiotropic leaf growth effects. Since PIF4 and TCP4 physically interact with each other, additional signaling components downstream to PIF4 and TCP4 together remain to be explored in response to environmental cues.

## Conclusions

The interplay between environmental signaling and leaf developmental pathway is understudied at the genetic level than either of the processes itself. Here, we showed the convergence of a developmental pathway and environmental signaling on the core cell cycle machinery. In our opinion, the discovery of high-temperature-mediated transcriptional induction of *KRP1*, PIF4 dependency on TCP4, and its involvement in leaf size regulation are new. The mechanism has a potential to be explored further to influence leaf growth plasticity in response to the temperature that can be targeted to optimize leaf growth under temperature changes.

## Materials and methods

### Plant Material and Growth Conditions

Arabidopsis plants (Columbia ecotype) were grown on half-strength Murashige and Skoog (Duchefa) plates at 21 ºC or 28 ºC with 100 μmolm^-2^s^-1^ light intensity under 16/8hrs light/dark conditions. The seeds of *pCyclinB1:GUS* (CS68143), *pPIF4:GUS* (CS69169), *pif4-2* (CS66043), and *pif7-1* (CS68809) were obtained from the Arabidopsis Biological Resource Center at Ohio State University. The mutants *tcp4-2* (GABI_363H08), *jaw1-D, krp1, krp4/6/7*, and the transgenic lines, 35S::TCP-3F6H and *pTCP4:GUS*, were described previously (Jakoby *et al*., 2006; Sarvepalli & Nath, 2011; Challa *et al*., 2016; Sizani *et al*., 2019; Kubota *et al*.,2017). The homozygous seeds of mutants *pif4tcp4* and *pif4krp1* were obtained by crossing. Primer details for genotyping of the mutants are available in Supplemental file S2.

### Microscopic Morphological Analysis and Cellular Analysis

Rosette pictures from the top were taken from three replicate media plates using a Canon EOS 700D camera. Images were converted to black and white images after color threshold changes and the rosette area was measured (as a sum of the area of all canopy leaves and their petioles) using ImageJ v 1.48 (http://rsbweb.nih.gov/ij/). Leaves were fixed overnight with 3:1 ethanol:acetic acid, followed by overnight clearing in 100% lactic acid. Cleared leaves were then pictured under a Nikon AZ-100 microscope equipped with a Nikon DS-Ri1 digital camera. Leaves with an area close to the average were chosen for drawing cells in ImageJ using a drawing tablet (Wacom) and analyzed as per Andriankaja et al. (2012).

### GUS assay

Whole plants were harvested at 10-12 DAS in GUS solution (Phosphate buffer (pH 7), 0.1% Triton X-100, 0.5 mM K4[Fe(CN)6].3H20, 0.5 mM K3[Fe(CN)6]) with X-gluc (958 μM X-gluc dissolved in 1% DMSO) and kept at 37 ºC for required time. Samples were fixed with ethanol:acetic acid (3:1) and cleared with 100% lactic acid before mounting on slides. The quantification of GUS staining was done using 4-methylumbelliferyl ß-D-glucuronide (4-MUG; Sigma) as a substrate with excitation at 365 nm and emission at 455 nm read using a plate reader POLARstar Omega (BMG Labtech) as previously described (Blazquez, 2007).

### Expression Profiling

RNA was extracted from proliferating leaves at 10-12 DAS using QIAzol lysis reagent (Qiagen) according to the manufacturer’s protocol. 1 μg of total RNA was quantified using a nanodrop spectrophotometer Nanovue (GE Healthcare) and used for making cDNA using Verso cDNA synthesis kit (Thermo Fischer Scientific). qPCR was performed using gene-specific primers using PowerUp SYBR Green master mix (Applied Biosystems) and Bio-Rad CFX9 connect thermocycler (Bio-Rad). Primers’ details can be found in Supplemental file S2. This was done as three biological and technical repetitions using ACT2 and TUB2 as the reference genes. The transcript levels were estimated using 2^-(dCT)^ method (Livak and Schmittgen, 2001; Schmittgen and Livak, 2008).

### Confocal Microscopy

Seven-day-old seedlings of *pPIF4:PIF4-GFP* were grown at 21 ºC were shifted to 21 ºC in dark or 28 ºC in dark for 4hours before taking images using Leica TCS SP5 confocal laser scanning microscope (Leica Microsystems). DAPI (Invitrogen) images were taken using 6-day old young seedling using Leica TCS SP8 confocal laser scanning microscope (Leica Microsystems).

### EdU fluorescence

Control and high temperature-grown 5-day old seedlings were harvested and incubated in liquid MS (half strength) containing 10nM EdU (Click-iT™ EdU Cell Proliferation Kit) for 30min in dark at room temperature. Seedlings were further incubated and washed with 1/2MS followed by washing in PBS buffer. As per the manufacturer’s protocol, seedlings were then incubated in a Click-iT reaction cocktail for 30min in dark followed by washes in PBS buffer. Fluorescence was detected using Alexa Fluor 488 settings with Leica TCS SP8 confocal laser scanning microscope (Leica Microsystems).

### Yeast Two-Hybrid Assays

The yeast two-hybrid assay was performed using the GAL4-based two-hybrid system (Clonetech, Mountain View, CA, USA). To check the interaction of PIF4 with TCP4, the full-length coding sequence of TCP4 with stop codon was amplified and cloned in pENTR-D-TOPO vector (Invitrogen). The prey construct i.e., TCP4-AD was generated *via* LR reaction between TCP4-TOPO and pGADT7g. Since we found that PIF4 can activate the reporter in yeasts, we excised 53 amino acids from the N-terminus of PIF4 to generate bait constructs. PIF4Δ53 was amplified from the cDNA using primers PIF4Δ53-CDS-F and PIF4-CDS-SC-R and cloned into the vector pENTR-D-TOPO to generate PIF4Δ53-TOPO. Further, the bait construct DBD-PIF4Δ53 was generated using LR reaction between PIF4Δ53-TOPO and pGBKT7g. Interaction between p53 (pGBKT7-p53) and large T-antigen (pGADT7-T) was used as a positive control and negative control was performed using pGBKT7-Lam and pGADT7-T. The interaction was checked between the bait construct DBD-PIF4Δ53 with each prey construct (TCP4-AD) or empty pGADT7g after co-transforming in yeast Y2H gold strain by using the EZ-Yeast™ Transformation Kit (MP Biomedicals) according to the manufacturer’s protocol.

### Bimolecular Fluorescence Complementation (BiFC) assays

For the BiFC assay, the full-length coding sequence of PIF4 (without stop codon) and TCP class II proteins (TCP2, TCP3, TCP4, TCP10, and TCP24) with stop codon were amplified and cloned into the vector pENTR-D-TOPO (Invitrogen). Further, using Gateway™ LR Clonase™ II Enzyme mix (Invitrogen), PIF4 CDS was cloned into pSITEcEYFP-N1 (CD3-1651) to generate PIF4-cYFP and TCPs CDS were cloned into pSITE-nEYFP-C1 (CD3-1648) to generate nYFP-TCPs constructs (Primer for each construct can be found in Supplemental file S2). These constructs were then transformed into Agrobacterium strain, EHA105. Agrobacterium cells harboring our constructs (PIF4-cYFP, nYFP-TCPs) and empty vectors were incubated in infiltration buffer (10 mM MES pH 5.6, 10 mM MgCl_2_, 200 μM acetosyringone) for 4 hours at 28 ºC. For co-infiltration, an equal volume of appropriate Agrobacterium cells were mixed and were co-infiltrated into the leaves of *Nicotiana benthamiana*. The infiltrated *N. benthamiana* plants were kept for overnight incubation in dark followed by light for 48 hours. YFP fluorescence signified a positive interaction. NLS-RFP was used as a nuclear marker. The YFP and RFP fluorescence were detected using Leica SP5 confocal laser scanning microscope (Leica Microsystems).

### Luciferase transactivation assay And Dual-Luciferase assay

For luciferase assay, the 1467 bp promoter region of KRP1 was amplified and cloned in the pENTR-D-TOPO vector (pKRP1-TOPO). Using Gateway™ LR Clonase™ II Enzyme mix (Invitrogen) and pKRP1-TOPO construct, pKRP1:LUC was generated using Gateway binary vector pGWB435:LUC. Full-length CDS of transcription factors, PIF4 and TCP4 were cloned under the control of constitutive 35S promoter into plant Gateway binary vector pGWB402, to generate 35S:PIF4 and 35S:TCP4, respectively. All these constructs and empty vectors were introduced into *A. tumefaciens* strain, EHA105. Agrobacterium cells containing 35S:PIF4+pKRP1:LUC and 35S:TCP4+pKRP1:LUC constructs were co-injected into *N. benthamiana* leaves. After 48 hours, the transformed leaves were sprayed with 100 mM luciferin, kept in dark for 3-5 minutes, and luminescence was detected using low light cooled CCD imaging apparatus (Bio-Rad). Further, to quantify luciferase activity under pKRP1, the same promoter region was used to generate pKRP1:LUC construct using gateway vector p635nRRF containing 35S:REN. The expression of REN was used as an internal control. LUC/REN ratio represented the relative activity of the *KRP1* promoter. All the mixed cultures (a. 35S:PIF4+pKRP1:LUC, b. 35S:TCP4+pKRP1:LUC, c. 35S:PIF4+35S:TCP4+pKRP1:LUC) were co-infiltrated into *N. benthamiana* leaves. Firefly luciferase and Renilla luciferase activities were assayed in extracts from leaf disc (1cm in diameter) using Dual-Luciferase Assay System E1910 (Promega) according to manufacturer’s protocol. The quantification was done using luminometer POLARstar Omega (BMG Labtech).

### Split Luciferase assay

PIF4 and TCP4 CDS were amplified using CDS primers with *kpn1* and *Sal1* restriction site and cloned into pJET1.2 vector (Thermo Fischer Scientific). Following restriction digestion, the products into the destination vectors pCAMBIA-cLUC (CD3-1700) and pCAMBIA-nLUC (CD3-1699), respectively, and co-infiltrated in *N. benthamiana* before imaging using Luciferase assay system E1500 (Promega) as mentioned above.

### Chromatin Immunoprecipitation

Arabidopsis seedlings were collected in dark at 12DAS grown at 21 ºC or shifted to 28 ºC for 8hours before harvesting them in crosslinking buffer containing formaldehyde. Chromatin Immunoprecipitation (ChIP) was performed using antiPIF4 antibody (AS16 3157; Agrisera) or Anti-FLAG(R) M2-Antibody (F1804; Sigma) as described in Saleh et al. (2008). 2μl chromatin was used for amplification using 40 cycles for qPCR analysis PowerUp SYBR Green master mix (Applied Biosystems) and CFXconnect thermocycler (Bio-Rad). Details for primers used in ChIP can be found in Supplemental file S2.

### Western Blot

8-day old seedlings were harvested at *Zt* 5 (*Zeitgeber time* 5) for control and high-temperature samples (treated between *Zt*1-*Zt*5). Protein from 200mg tissue was isolated using YODA-buffer (50 mM Tris pH7.5; 150 mM NaCl; 1 mM EDTA; 10% glycerol; 5 mM DTT; 1% protease inhibitor cocktail (Sigma); 10 μM MG132; 0.1% Triton-X100). Protein electrophoresis and western blot were performed (Bolt mini blot module, Invitrogen) using antiPIF4 antibody (AS16 3157; Agrisera) and Anti-RbcL antibody (AS03 037A; Agrisera), and detected using ECL chemiluminescence kit (BioRad). Imaging was done using Hi-sensitivity chemiluminescence settings under ChemiDoc XRS+ (BioRad).

### Quantification and statistical analysis

Quantitative data from all experiments were compiled and analyzed in GraphPad Prism v.8.3.2. One-way ANOVA and post hoc Tukey test were done using SPSS. Student’s t-tests were done using GraphPad. Conditions of normality of distribution and homogeneity of variance were checked and met. Details of the statistical tests applied, including the statistical methods, the number of replicates, mean, and error bar details, and significances are indicated in the relevant figure legends.

## Acknowledgments

We are grateful to Gerrit Beemster for sharing with us the seeds of *krp1*, and *krp4/6/7*, Christian Fankhauser for *pif1345*, Utpal Nath for *tcp4-2* and *pTCP4:GUS* lines, On Sun Lau for *pPIF4:PIF4-GFP*, and to Takato Imaizumi for the 35S::TCP-3F6H line. We thank Praveen Verma for providing the p635nRRF vector for dual luciferase assay. K.S. acknowledge her DBT-RA and SERB-NPDF fellowship, and A.D. acknowledge her UGC-JRF fellowship. We thank the Confocal facility and other Central Instrumentation Facility (CIF) at NIPGR for their support.

## Author Contributions

A.R. and K.S. conceived the project and designed the experiments. K.S. and A.D. performed the experiments. K.S. analyzed the data and wrote the manuscript with contributions from A.R. and A.D. A.R. supervised the project, provided scientific discussions, and agrees to serve as the author responsible for contact and ensures communication.

## Funding Information

This work was supported by Ramalingaswamy Re-entry Fellowship (BT/RLF/Re-entry/05/2013) from the Department of Biotechnology, Ministry of Science and Technology, India, and SERB-Core Research Grant (CRG/2020/002179) and SERB-NPDF scheme (PDF/2018/002387) from Science and Engineering Research Board, India.

## Supplemental Data

**Supplemental Figure S1**. Leaf phenotype in control and high-temperature grown Arabidopsis plants.

**Supplemental Figure S2**. Phenotyping of the first pair of leaves under high temperature.

**Supplemental Figure S3**. Quantification of the GUS activity in *pCyclinB1;1:GUS* reporter line and fluorescent intensity in EdU treated Col-0 leaves.

**Supplemental Figure S4**. Proliferating leaves of Col-0 seedlings stained with DAPI.

**Supplemental Figure S5**. PIFs control temperature-mediated leaf architectural changes.

**Supplemental Figure S6**. Leaf features in control and high temperature grown Arabidopsis mutants.

**Supplemental Figure S7**. Quantification of the GUS activity in control and high temperature treated leaves of the *promoter:GUS* reporter lines of *TCP4*.

**Supplemental Figure S8**. Quantification of rosette area in *PIF4-* and *TCP4-* related mutants under control and high temperature.

**Supplemental Figure S9**. Quantification of rosette area in *PIF4-* and *KRP1-* related mutants under control and high temperature.

**Supplemental Figure S10**. BiFC assays in *N. benthamiana* showing protein-protein interaction between PIF4 and class II TCP members.

**Supplemental File S1**. Literature curated leaf development-related genes that were among DEGs under high-temperature conditions in Arabidopsis.

**Supplemental File S2**. List of primers used in the study.

## Notes

### Competing Interest Statement

The authors have declared no competing interest.

