## Supplemental Figure for "High temperature limits leaf size via direct control of cell cycle by coordinated functions of PIF4 and TCP4"

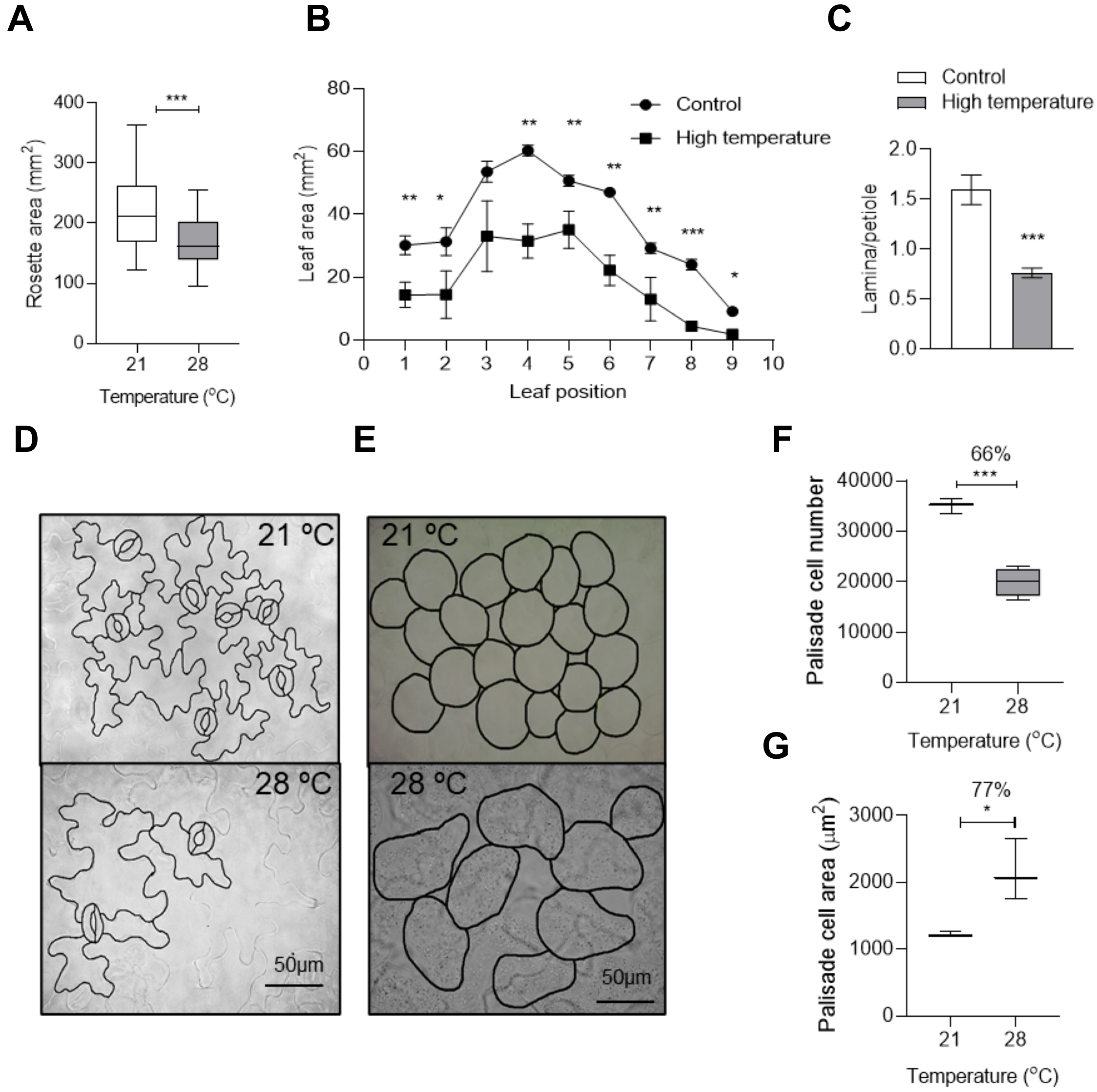


**Supplemental Figure S1.** Leaf phenotype in control and high-temperature grown Arabidopsis plants. (A) Quantification of rosette area of 21 ºC and 28 ºC grown Arabidopsis Col-0 wild type plants at 20 DAS (n=80-100). (B) Leaf series showing leaf area difference in Col-0 plant grown at 21 ºC and 28 ºC at 20 DAS (n=4). (C) Lamina/petiole ratio of leaf four in Col-0 at 20 DAS (n=8). (D, E) Representative images showing (D) abaxial epidermal cells and (E) palisade mesophyll cells in leaf four under control (top) and high temperature (bottom) conditions at 20DAS. Fully visible cells were hand drawn using ImageJ. Scale, 50µm. (F, G) Quantification of average palisade cell number (F) and average palisade cell area (G) in leaf four under control and high temperature conditions (n=3). Values represent mean±SE. Stars indicate **P* value <0.05, ***P* value<0.01, ****P* value <0.001 using the student’s t test. Percentage denote percent change between control and high temperature condition (F, G).

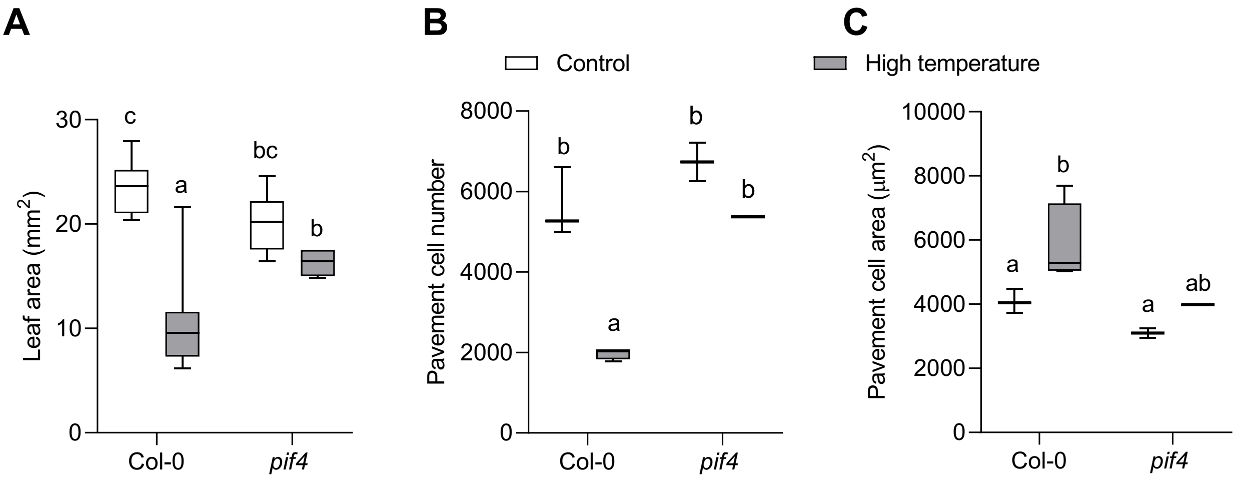


**Supplemental Figure S2.** Phenotyping of the first pair of leaves under high temperature. (A-C) Quantification of leaf area (n=14-18, A), Average pavement cell number per leaf in the adaxial side (n=3, B), and Average pavement cell area in the first pair of leaves (C) under 21 ºC and 28 ºC at 20 DAS. DAS, days after stratification. Box plot extends from $25^{th}$ to $75^{th}$ percentiles, where line represents the median and whiskers minimum and maximum value. Letters above the bars indicate statistically significant differences between genotypes or treatments (one way ANOVA followed by post hoc Tukey's test, $P < 0.05$).

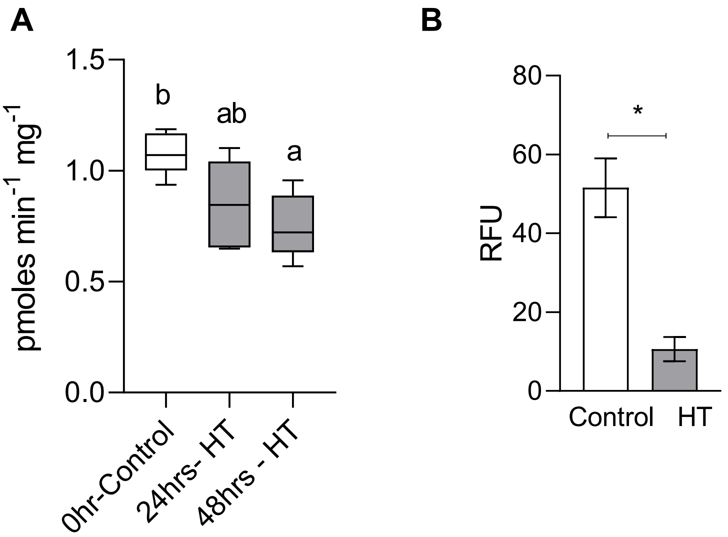


**Supplemental Figure S3.** (A) Quantification of the GUS activity in *pCyclinB1;1:GUS* reporter line under control and high-temperature conditions. Values represent mean ± SE (n=6). Box plot extends from 25th to 75th percentiles, where line represents the median and whiskers min. and max value. Letters above the bars indicate statistically significant difference between treatments (one way ANOVA followed by post hoc Tukey's test, $P < 0.05$). (B) Quantification of fluorescent intensity in EdU treated Col-0 leaves under control and high-temperature conditions. RFU= Relative fluorescent unit (arbitrary fluorescent unit). Values represent mean ± SE (n=3). Stars indicate student's t- test **P* value <0.05. HT- High temperature

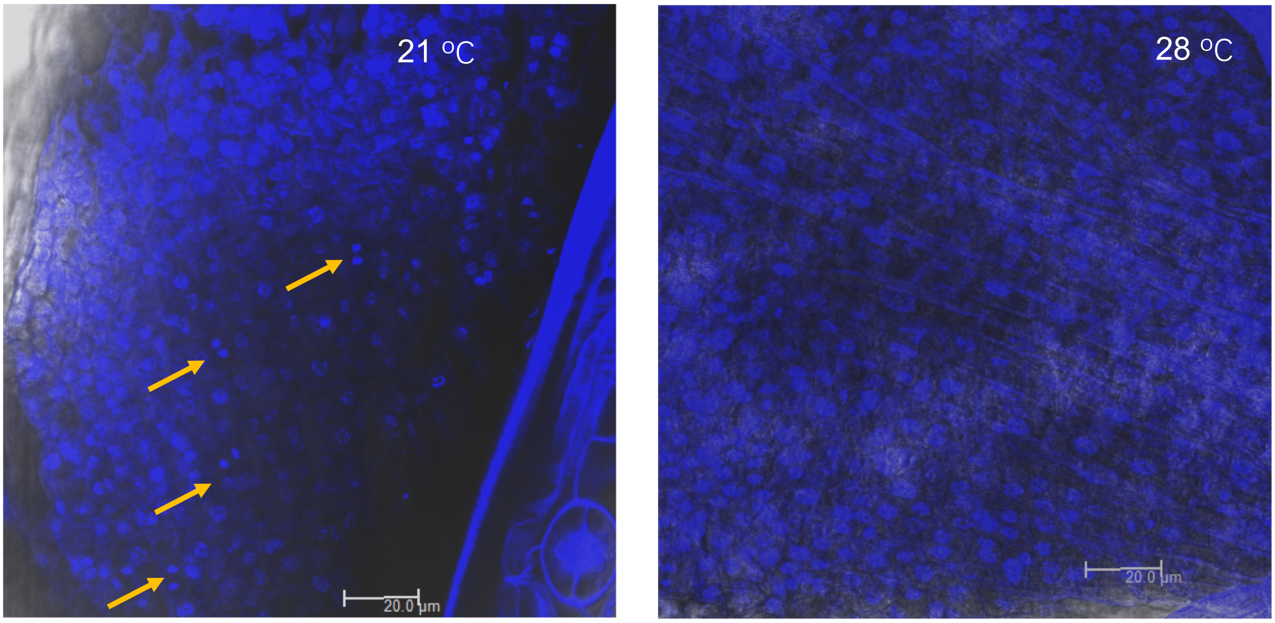


**Supplemental Figure S4**. Proliferating leaves of Col-0 seedlings stained with DAPI (4′,6-diamidino-2-phenylindole) to visualize mitotic figures as a marker for cells undergoing division under control and high temperature grown seedlings. Arrows indicate cells in anaphase. Scale, 20µm.

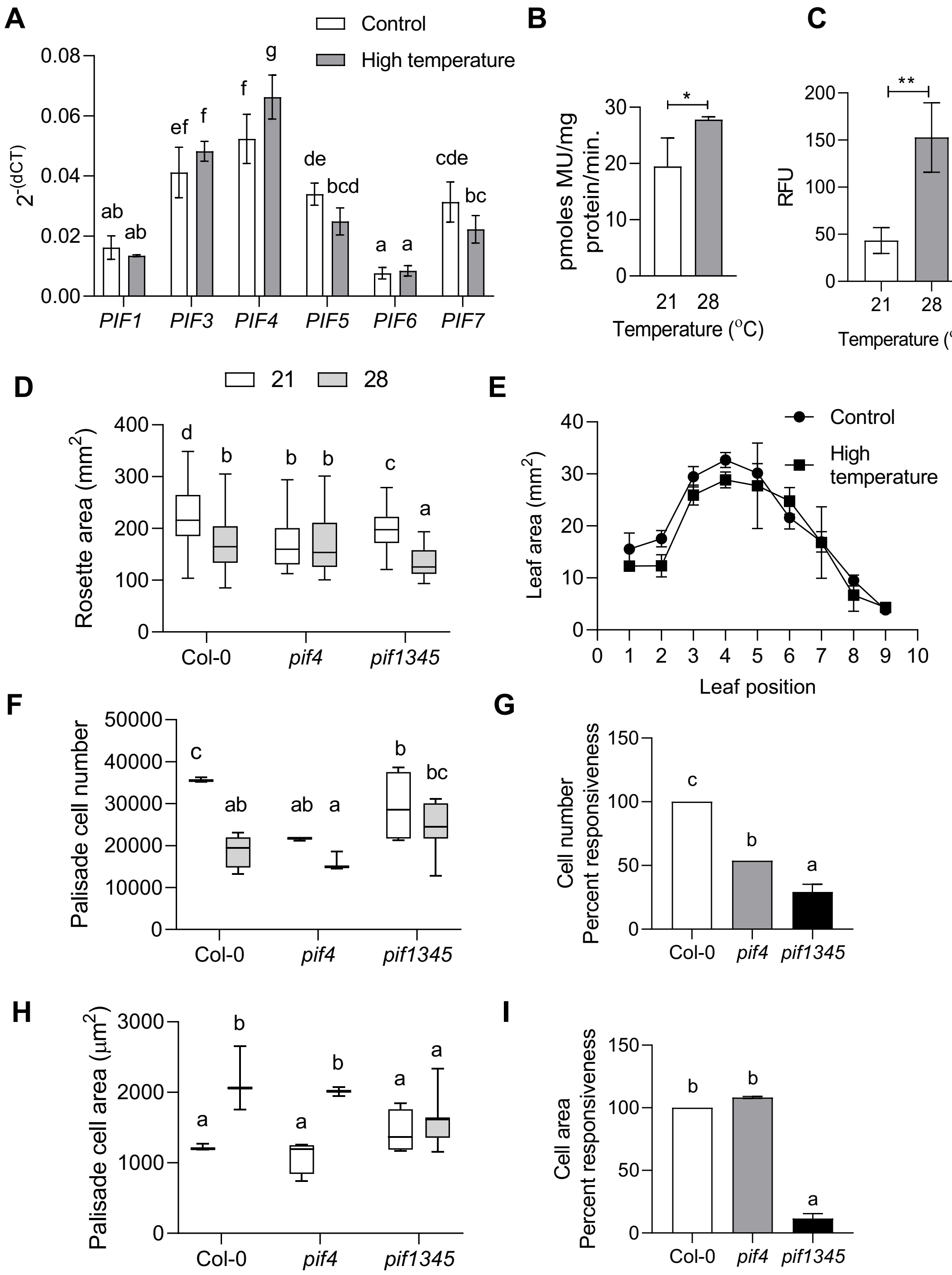


**Supplemental Figure S5.** PIFs control temperature-mediated leaf architectural changes. (A) Expression level of *PIF*s relative to an endogenous control, *ACTIN2*, in control and 8hr high-temperature treated young proliferating leaves of Col-0 plotted as $2^{-(dCT)}$. Values represent mean ± SE (n=6). (B) Quantification of the GUS activity in control and high temperature treated leaves of the *promoter:GUS* reporter lines of *PIF4*. Values represent mean ± SE (n=3). (C) Quantification of the fluorescence in the epidermal cells of a translation fusion reporter line *pPIF4:PIF4-YFP* under control and high temperature. Scale, 20μm. Values represent mean ± SE (n=4). (D) Quantification of rosettes area at 20DAS. (E) Leaf series showing leaf area of different rosette leaves of *pif4* mutant grown under control and high temperature conditions for 20days. Values represent mean ± SE (n=4). (F) Average palisade cell number in the abaxial side of leaf four (n=3). (G) Percent responsiveness of various genotypes to high temperature calculated with respect to the wild type Col-0 after setting the percent response of Col-0 to 100%. (H) Average palisade cell area in the abaxial side of leaf four (n=3). (I) Percent responsiveness of various genotypes to high temperature calculated with respect to the wild type Col-0 after setting the percent response of Col-0 to 100%. Box plot extends from 25th to 75th percentiles, where the line represents the median and whiskers minimum and maximum values. Letters above the bars indicate statistically significant differences between genotypes or treatments (one way ANOVA followed by post hoc Tukey's test, $P < 0.05$). Stars indicate students' t-test *$P$ value <0.05, **$P$ value <0.01.

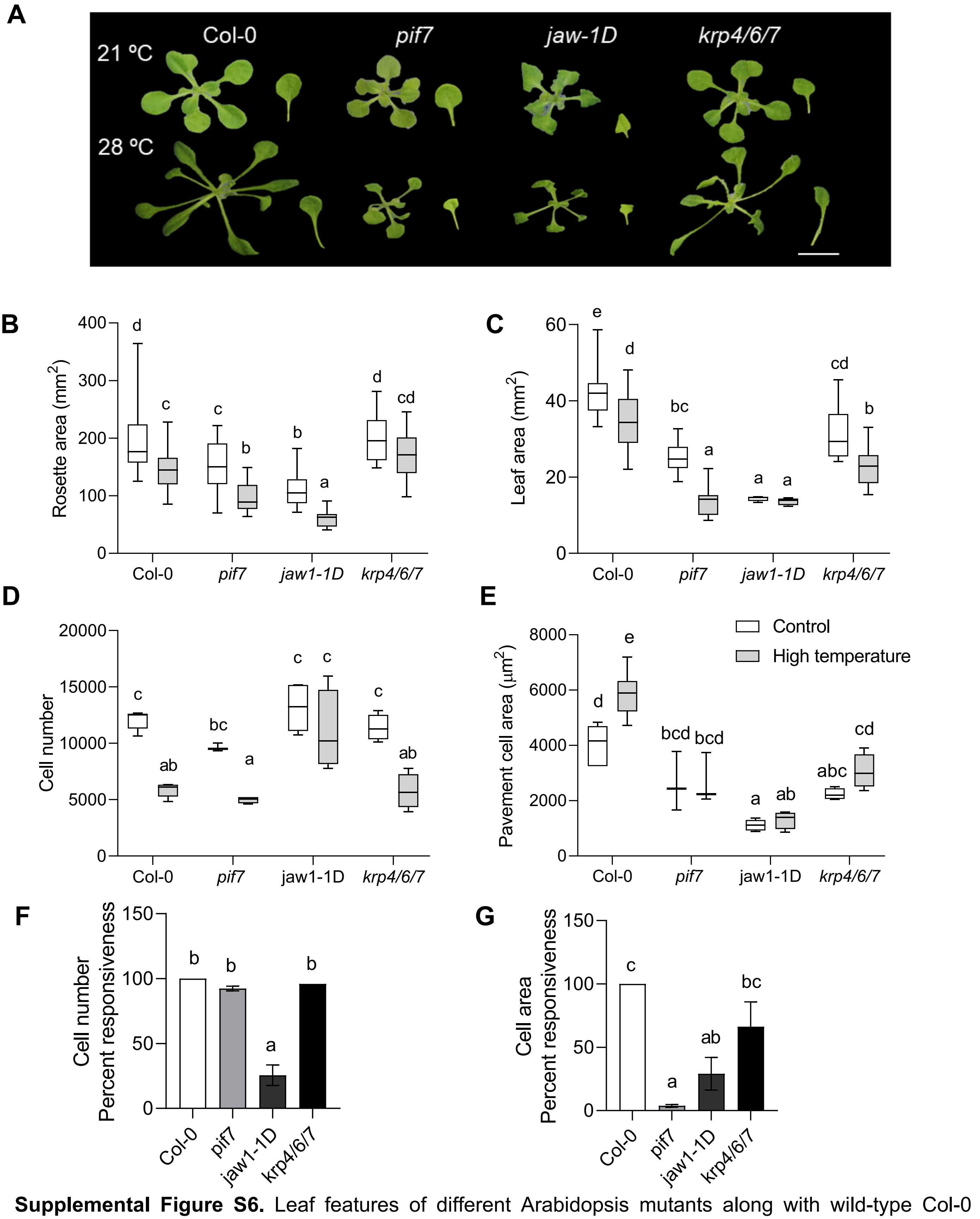


**Supplemental Figure S6.** Leaf features of different Arabidopsis mutants along with wild-type Col-0 under control and high temperature conditions. (A) Representative rosette and leaf four images of Col-0 and various mutants (as indicated in the figure) in 21 $^0$C and 28 $^0$C grown plants at 20 DAS. (B-E) Quantification of rosette area (n>50, B), leaf four area (n=15-20, C), average pavement cell number per leaf in the adaxial side (n=5-6, D), and average pavement cell area (E) in 21 $^o$C and 28 $^o$C grown plant at 20 DAS (n=5-6). (F, G) Percent responsiveness of different mutants to high temperature calculated with respect to the wild type Col-0 after setting the percent response of Col-0 to 100%. Box plot extends from 25th to 75th percentiles, where line represents the median and whiskers min. and max value. Letters above the bars indicate statistically significant differences between genotypes or treatments (one way ANOVA followed by post hoc Tukey's test, $P < 0.05$).

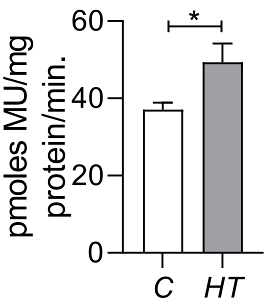


**Supplemental Figure S7.** Quantification of the GUS activity in control and high temperature treated leaves of the *promoter:GUS* reporter lines of *TCP4*. Values represent mean ± SE (n=3). Star indicate students' t-test **P* value < 0.05. C, control; HT, high temperature.

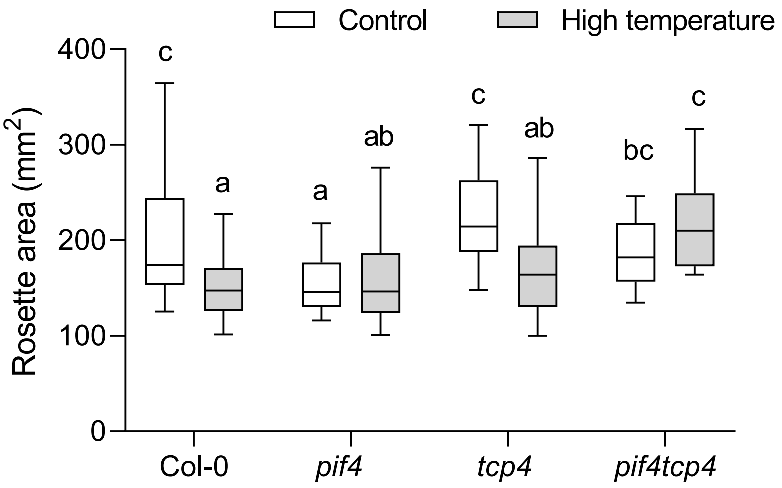


**Supplemental Figure S8.** Quantification of rosette area in *PIF4*- and *TCP4*- related mutants under control and high temperature (n=45-55). Box plot extends from 25th to 75th percentiles, where the line represents the median and whiskers minimum and maximum values. Letters above the bars indicate statistically significant differences between genotypes or treatments (one way ANOVA followed by post hoc Tukey's test, $P < 0.05$).

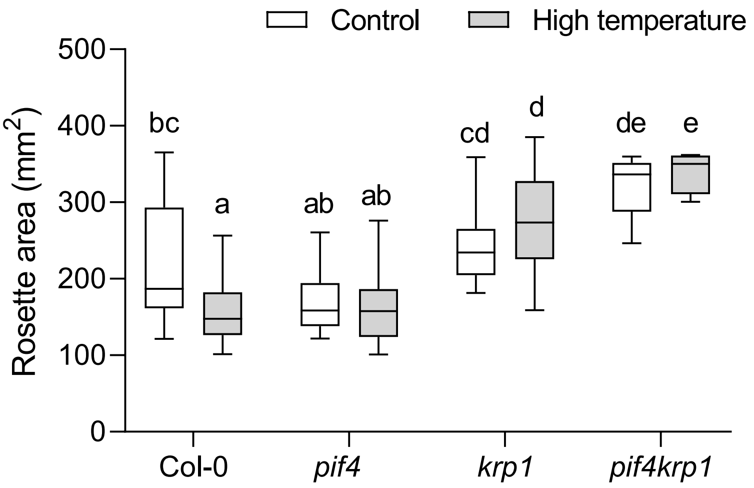


**Supplemental Figure S9.** Quantification of rosette area in *PIF4*- and *KRP1*- related mutants under control and high temperature (n=30-38). Box plot extends from 25th to 75th percentiles, where the line represents the median and whiskers minimum and maximum values. Letters above the bars indicate statistically significant differences between genotypes or treatments (one way ANOVA followed by post hoc Tukey's test, $P < 0.05$).

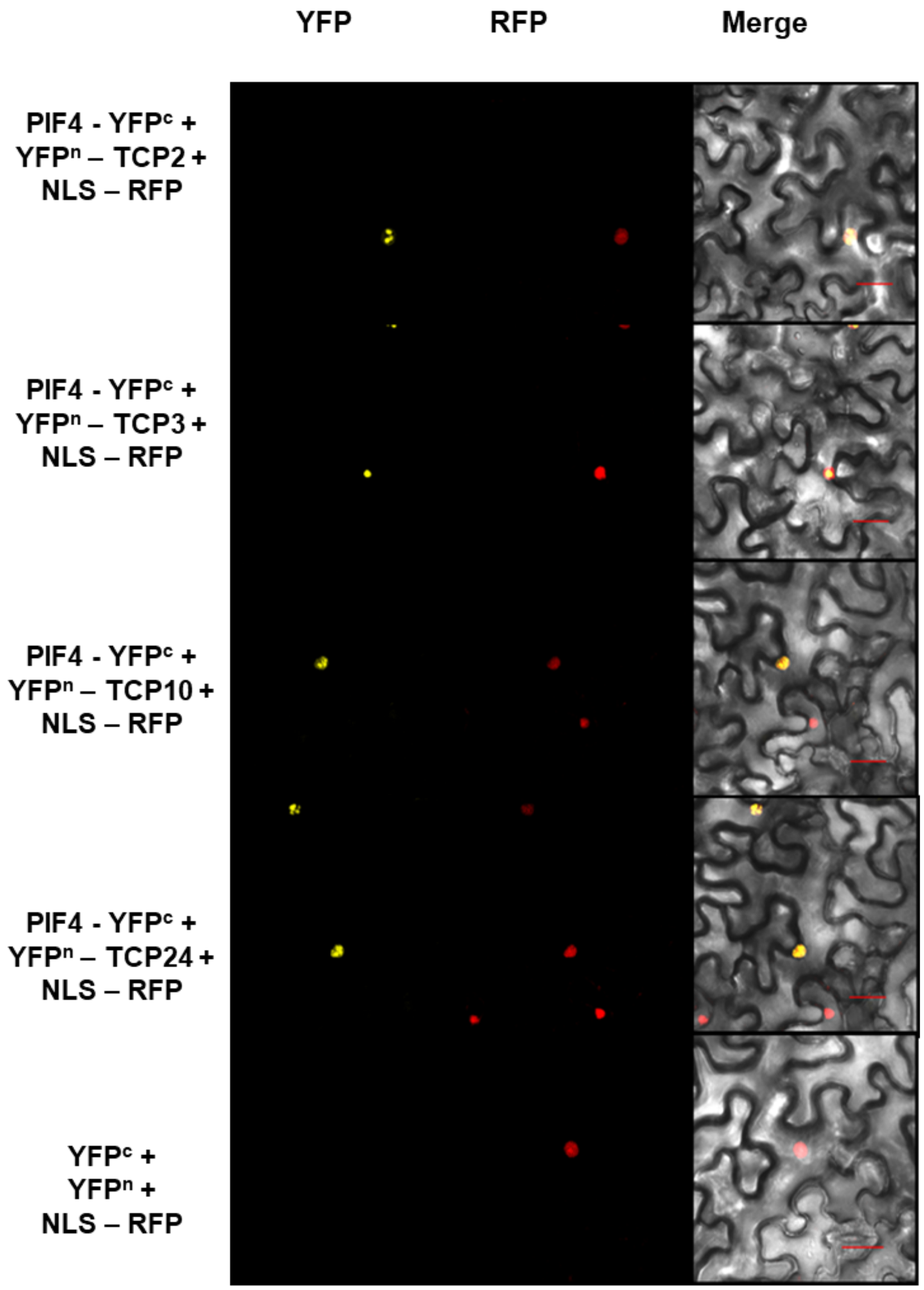


**Supplemental Figure S10.** BiFC assays in *N. benthamiana* showing protein-protein interaction between PIF4 and class II TCP members i.e. TCP2, TCP3, TCP10 and TCP24 from top to bottom. Empty vector (bottom) was used as negative control. NLS-RFP was used as a nuclear marker. (Scale, 30 µm).
